# DUSP12 regulates NAT10-mediated RNA acetylation to modulate DNA repair and therapeutic response in hepatocellular carcinoma

**DOI:** 10.64898/2026.01.10.698822

**Authors:** Viktor Kalbermatter Boell, Diana Reis Della Corte Guimarães Pacheco, Yuli Thamires Magalhães, Isabeli Yumi Araújo Osawa, Nicolas Carlos Hoch, Fábio Luís Forti

**Author notes:** **Corresponding author:** Fabio Luis Forti, PhD, Av. Prof. Lineu Prestes, 748 - Bl.09i, Sl.922, CEP: 05508-900 – Cidade Universitária, São Paulo-SP, Brazil /Fax: 55-11-3091-2186. **First author:** Viktor Kalbermatter Boell.

## Abstract

Hepatocellular carcinoma (HCC), the most common type of primary liver cancer arising from hepatocytes, is an aggressive hepatic malignancy with limited therapeutic options and poor prognosis. Chemotherapy remains an important treatment for advanced disease, though the mechanisms influencing drug sensitivity remain elusive. This study investigates the role of dual-specificity phosphatase 12 (DUSP12) and its interaction with the nucleolar protein NAT10 in the hepatic cellular response to genotoxic stress. We demonstrate that doxorubicin (DX) induces superior cytotoxicity over cisplatin in hepatocellular carcinoma models, associated with a stronger DNA damage response (DDR), nucleolar stress, and delocalization of NAT10 from the nucleolus to the nucleoplasm, where it colocalized with DUSP12. Genetic ablation of DUSP12 sensitized cells to DX, increasing DNA damage markers (p53, p-p53(Ser15), γH2AX(Ser139)) and delaying the repair of DNA strand breaks. DUSP12 knockout also caused redistribution of nucleolar proteins NAT10 and TCOF1. Pharmacological inhibition of NAT10 in DUSP12-deficient cells further enhanced DX sensitivity, revealing a synthetic-lethal interaction. We identified a direct association between NAT10 and DUSP12’s functional domains, with evidence indicating NAT10 is a DUSP12 substrate. Consequently, DUSP12 knockout elevated NAT10 phosphotyrosine levels and reduced ac4C RNA acetylation, indicating functional impairment of NAT10. Corroborating these findings, patient data showed frequent DUSP12 amplification in HCC, correlating with poor survival and enrichment in DDR and ribosome biogenesis pathways. Our results establish the DUSP12-NAT10-ac4C axis as a molecular link between the DDR and nucleolar stress, highlighting previously unrecognized therapeutic vulnerability in HCC.

## Introduction

Hepatocellular carcinoma (HCC) is the most prevalent primary liver malignancy, accounting for over 906,000 diagnosed cases globally in 2020. HCC is the third leading cause of cancer-related mortality worldwide, with a relative 5-year survival rate of approximately 18% [1]. Its aggressive nature and poor prognosis are primarily attributed to late-stage diagnosis and the limited efficacy of available therapeutic options, which depend largely on combinations of tyrosine kinase inhibitors (TKIs) and immune checkpoint inhibitors [2]. Chemotherapy remains a key therapeutic strategy for HCC, including its use in transarterial chemoembolization (TACE) with agents such as doxorubicin and cisplatin, which exert anticancer effects by inducing DNA damage and activating the DNA damage response (DDR) – a highly coordinated signaling network that detects DNA lesions, induces cell cycle arrest, and promotes repair to maintain genomic stability [3]. Notably, evidence suggest a functional interplay between the DDR and ribosome biogenesis (RB) pathways, highlighting their potential relevance in cancer pathophysiology.

RB is a complex and energy-intensive process involving the synthesis, processing, and assembly of ribosomal RNA (rRNA) and ribosomal proteins [4]. The nucleolus, beyond its traditional role in ribosome production, acts as a central hub for sensing cellular stress, including DNA damage [5,6]. Disruption of nucleolar function, often termed “nucleolar stress”, activates DDR by triggering the release of key nucleolar components, such as ribosomal proteins, which can stabilize p53 and other DDR effectors [7,8]. Conversely, several DNA damage response-related proteins can impair RB [9–12]. In cancer, the interplay between DDR and RB is particularly significant since many oncogenic signaling pathways involve overactivation of rRNA synthesis leading to uncontrolled cell proliferation [13,14]. However, these changes also increase sensitivity to nucleolar stress and DNA damage, creating potential therapeutic vulnerabilities. For instance, targeting RB via nucleolar disruption has been shown to sensitize chemoresistant cells to genotoxic therapies [15–17].

The dual-specificity phosphatase 12 (DUSP12), a relatively unexplored phosphatase in humans, seems to be implicated in both genotoxic stress response and nucleolar function. DUSP12 displays a distinctive and conserved zinc-binding domain which points to the possibility of interactions with nucleic acids and ribonucleoproteins [18,19]. Like its ortholog Yvh1 in yeast, DUSP12 is now recognized for its involvement in late stages of ribosome biogenesis [20–23]. Moreover, through dephosphorylation of its sole known substrate, ASK1, and subsequent attenuation of the JNK/p38 MAPK pathway, DUSP12 expression and activity are critical for regulating hepatic lipid metabolism and ensuring cell survival under conditions such as diabetic cardiomyopathy, hepatic and neuronal ischemia-reperfusion [24–27]. Interestingly, DUSP12 physically interacts with nucleolar proteins under genotoxic stress conditions, such as N-acetyltransferase 10 (NAT10), which is a regulator of ribosomal RNA transcription and processing [28]. NAT10 is the only enzyme identified in humans as a writer of N4-acetylcytidine (ac4C) across a broad range of RNAs: the acetylation of mRNA and tRNA enhances translational efficiency and fidelity [29,30]. In addition to acetylating 18S rRNA and promoting its maturation, NAT10 contributes to RB by acetylating UBF, thereby facilitating RNA Pol I recruitment to rDNA promoter regions [31,32]. Through acetylation of protein substrates, NAT10 is also implicated in cellular processes such as cell division, autophagy, chromatin remodeling, and DNA damage response [33–36]. Interestingly, NAT10 is upregulated and related to progression, metastasis, and chemoresistance of HCC [37–39].

The interactions between DUSP12 and NAT10 suggest this axis may serve as a molecular bridge linking DNA damage and nucleolar stress pathways acting over nucleolar proteins. Thus, this study aimed to investigate this interplay in liver cells, more specifically HCC, focusing on the involvement of the DUSP12-NAT10 axis in these processes. Two genotoxic agents used in chemotherapy were used to sensitize 2D and 3D cultures of two different liver cells, which had previously DUSP12 knocked out. Our findings showed that loss of DUSP12, associated or not with NAT10 inhibition, caused sensitization of both cells to stress with impairment of DDR, DNA repair, and nucleolar reorganization.

## Materials and Methods

### Cell culture and treatments

Hepatic cell lines HepaRG and HuH-7 were maintained in DMEM supplemented with 10% FBS, ampicillin (25 µg/mL), and streptomycin (100 µg/mL) at 37 °C with 5% CO₂. Doxorubicin (Sigma), actinomycin D (Invitrogen), and remodelin (MedChem Express) were prepared as 10 mM stock solutions in DMSO, and cisplatin (Sigma) was prepared as 10 mM in 0.9% NaCl.

### CRISPR/Cas9-mediated knockout of DUSP12

DUSP12 knockout was generated using sgRNAs cloned into the eSpCas9(1.1) vector (AddGene #71814) (sequences in Supplementary Table 1). Cells were cotransfected with pEGFP-C1, selected with G418 (1 µg/mL, 21 days), and clones were isolated by serial dilution.

### Cell viability and proliferation assays

Cell viability and proliferation in monolayer cultures were assessed using the MTT (Invitrogen) reduction and the crystal violet incorporation assay, respectively. For cell viability, cells were incubated with 500 µg/mL MTT for 3 hours at 37°C, followed by solubilization of formazan in DMSO and absorbance measurement (570 nm). For cell proliferation, cells were stained with 0.5% crystal violet in 20% methanol for 10 minutes, followed by the incorporated crystal violet elution in 33% acetic acid and absorbance measurement (590 nm). IC_50_ values were calculated using non-linear regression with GraphPad Prism v.10.

### Spheroid formation and viability

Spheroids were generated via the liquid overlay method on 96-well plates coated with 1.5% agarose [40]. Morphometry was performed to evaluate spheroid size, using phase-contrast microscopy (Leica DMi1, 5X objective). Viable cell populations in intact spheroids were quantified with the acid phosphatase assay (APH), measuring absorbance of p-nitrophenolate at 405 nm [41].

### Immunofluorescence and confocal microscopy

Cells were seeded on glass coverslips, fixed with 4% PFA, and permeabilized with Triton X-100 on ice under optimized conditions: 0.5% for 5 min for HepaRG, or 1% for 10 min for HuH-7 and derived cell lines. Blocking was carried out in 3% BSA with 10% FBS, followed by primary antibodies (overnight, 4°C) and Alexa Fluor-conjugated secondary antibodies (1 h, room temperature) incubations. All antibodies used are listed in Supplementary Table 2. Nuclei were counterstained with DAPI, and slides were mounted with ProLong Diamond. Images were acquired using a Leica DMi8 fluorescence microscope, and confocal imaging was performed with a Leica SP8 STED FALCON system.

### Western blotting

Cell fractionation and total cellular lysates were obtained and quantified as previously described [42,43]. Equal amounts of protein (25-50 µg) were separated by 12% SDS-PAGE and transferred to nitrocellulose membranes (Millipore). Membranes were blocked in 3% BSA in TBST, followed by primary antibodies (overnight, 4°C) and Alexa Fluor-conjugated secondary antibodies (1 h, room temperature) incubations. All antibodies used are listed in Supplementary Table 3. Detection was performed with the Odyssey Infrared Imaging System. Quantification was performed in ImageJ by densitometry and normalized by the control condition.

### Cell cycle analysis

Exponentially growing cells were incubated with 10 µM BrdU (Invitrogen) for 30 min, followed by fixation and permeabilization as described previously. Acid-wash protocol was performed according to the manufacturer’s instructions. Blocking and immunodetection were performed as previously described using anti-BrdU and anti-geminin antibodies (Supplementary Table 2). Fluorescence microscopy images were acquired on a custom TissueFAXS i-Fluo system (TissueGnostics) mounted on a Zeiss AxioObserver 7 and an ORCA Flash 4.0 v3 camera (Hamamatsu). Image analysis was performed using StrataQuest software (TissueGnostics).

### Comet assay

Alkaline and neutral comet assays were performed to detect DNA fragmentation, as described previously [44] with modifications. Cells were lysed and deproteinized (2.5 M NaCl, 100 mM EDTA, 1% Triton X-100, 10% DMSO, 10 mM Tris, pH 10), DNA was denatured, and electrophoresis was carried out in alkaline buffer (300 mM NaOH, 1 mM EDTA, pH > 13) for 30 min or in neutral buffer (300 mM NaOAc, 100 mM EDTA, pH 8.3) for 1 h, both at 300 mA and 4°C. Slides were neutralized with 400 mM Tris (pH 7.4), fixed in ethanol, stained with SYBR Green II (Invitrogen), and analyzed using Comet Assay IV software (Instem).

### Detection of intracellular ROS levels

ROS levels were quantified using the CellROX Deep Red probe (Invitrogen) as described previously, according to the manufacturer’s instructions [42].

### RNA extraction and dot blot

Total RNA was isolated using TRIzol reagent (Invitrogen). RNA purity (A260/A280 = 1.8-2.1) was verified with a NanoDrop spectrophotometer, and integrity was assessed using the Agilent RNA 6000 Nano kit on a Bioanalyzer system. For ac4C detection, 500 ng of RNA was denatured at 65°C for 30 min in 1 mM EDTA (pH 8.0), 4% formaldehyde, and 10X SSC (5:2:3), then blotted onto Hybond N+ membranes using a Bio-dot apparatus (Bio-Rad) and UV-crosslinked (120 mJ/cm², 1 min). Membranes were stained with methylene blue (50 µg/mL in 5% acetic acid) as loading control. Blocking and immunodetection were performed as previously described using anti-ac4C antibody (Supplementary Table 3). Detection and quantification were performed as previously described.

### Pull-down and NAT10-immunoprecipitation assays

Plasmids to produce recombinant DUSP12 (full-length, N-terminal, and C-terminal domains) were kindly provided by Prof. Vacratsis [45]. Proteins were expressed in *E. coli* BL21-DE3 and purified from bacterial lysates using glutathione-agarose beads (Invitrogen). For pull-down assays, cell lysates were incubated with recombinant GST-tagged DUSP12 constructs at a 10:1 mass ratio, as previously described [46], in the presence or absence of remodelin and Na_3_VO_4_. Following incubation, the beads were directly applied to 12% SDS-PAGE for subsequent detection of NAT10. Normalization was performed using a GST-specific primary antibody (Supplementary Table 3) and was further adjusted based on the total lysate input to account for global protein expression under drug-treated conditions (doxorubicin and cisplatin) or in the spheroids culture model. For NAT10 immunoprecipitation, the Protein A/G Plus kit (Santa Cruz Biotechnology, sc-2003) was used according to the manufacturer’s instructions. Following incubation with cell lysates, detection was performed using a phosphotyrosine-specific primary antibody (Supplementary Table 3), and the signal was additionally normalized to the overall NAT10 expression levels in the corresponding lysate.

### Bioinformatics analysis

Liver hepatocellular carcinoma data from TCGA (PanCancer Atlas, n = 372) available on cBioPortal was utilized to analyze DUSP12 expression and genetic alterations in HCC [47]. Co-expressed genes (ρ > 0.3, n = 1189) were identified using the same database and subjected to gene ontology analysis on ShinyGO 0.81 with FDR cutoff =

0.05 [48].

### Statistical analysis

Experiments were conducted with at least three biological replicates, each with two to six technical replicates. Statistical significance was assessed using two-way ANOVA with post hoc tests (Tukey’s or Sidak’s) or one-way ANOVA for two-group comparisons, using GraphPad Prism 10. Statistical significance was denoted as p<0.05 (* or #), p<0.01 (** or ##), p<0.001 (*** or ###), p<0.0001 (**** or ####), and ns = not significant.

## Results

### Doxorubicin is slightly more cytotoxic than cisplatin in monolayers of hepatic cells

The human HCC cell line HuH-7 was treated with different genotoxic agents to assess whether cellular sensitivity depended on the type of DNA damage induced. Both cisplatin (CP) and doxorubicin (DX) reduced cell viability, but DX exerted a more pronounced effect on proliferation inhibition (Figure 1A–1D, Supplementary Table S4– S5). Similar results were obtained for the hepatic progenitor cell line HepaRG (Figure S1A–S1D). To mimic physiological conditions, both treatments were tested in spheroids. Standardization of cell number and culture duration enabled the generation of spheroids with physiologically relevant diameters (> 350 µm) in both HuH-7 (Figure S2A–S2B) and HepaRG lines (Figure S2D–S2E). Although spheroid diameter remained constant or even decreased under certain conditions, APH viability assays revealed an increase in the population of viable cells, suggesting that these changes reflect spheroid reorganization rather than inhibition of proliferation (Figure S2C, S2F). Compared to monolayers, HuH-7 spheroids were significantly less responsive to CP and DX treatments (Figure 1E–1F, Supplementary Table S6). DX treatment resulted in a 150-fold increase in IC_50_ for spheroids versus monolayers (IC_50_ monolayer, 72 h = 0.351 µM; IC_50_ spheroid, 72 h = 52.9 µM). Similar results were observed for HepaRG spheroids (Figure S1E–S1F).

**Figure 1.**
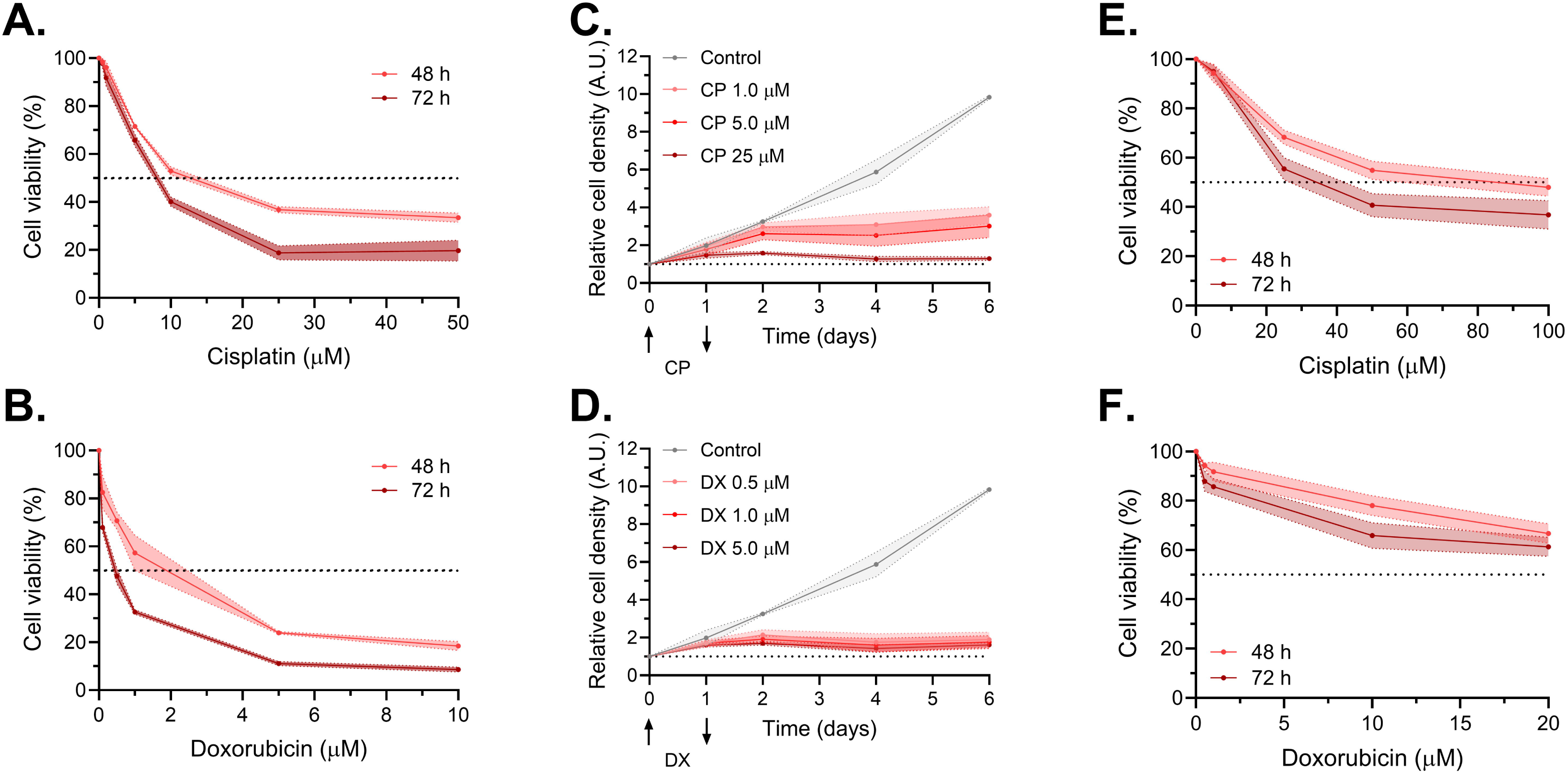
Cytotoxic effects of cisplatin and doxorubicin in monolayers and spheroids of HuH-7 cells. (A–B) Cell viability of HuH-7 monolayers treated with increasing concentrations of (A) cisplatin (CP) or (B) doxorubicin (DX) for 48 or 72 h. (C–D) Proliferation curves of HuH-7 monolayers treated with (C) CP or (D) DX for 24 h. (E–F) Cell viability of HuH-7 spheroids treated with increasing concentrations of (E) CP or (F) DX for 48 or 72 h. Data represent mean ± SD of at least three independent experiments.

### Doxorubicin induces stronger DDR activation, nucleolar stress features, NAT10 translocation, and enhanced colocalization with DUSP12

As evidenced by increased levels of phospho-p53 (Ser15) and quantification of γH2AX foci, DX induced a stronger and earlier DDR activation compared to CP in both HuH-7 (Figure 2A–2B, 2F–2G) and HepaRG cells (Figure S3A–S3B, S3F–S3G). The expression and phosphorylation of p53 varied according to the mutational status of the protein in each cell line (Figure 2A–2C, S3A–S3C). DUSP12 expression remained unchanged upon treatment, whereas DX induced a moderate reduction in NAT10 expression in both models (Figure 2A, 2D–2E, S3A, S3D–S3E). Subcellular distribution of DUSP12 between cytoplasmic and nuclear compartments was not affected by either treatment, as shown by immunofluorescence (Figure 2F, S3F) and fractionation assays (Figure S4). Interestingly, in HepaRG cells DUSP12 was predominantly nuclear, whereas this distribution pattern was lost in HuH-7 cells. In contrast, NAT10 localization was markedly altered in response to DX, with translocation from the nucleolus to the nucleoplasm (Figure 2F, S3F). This NAT10 redistribution was accompanied by alterations in the number of nucleoli per nucleus (Figure 2H, S3H), resembling nucleolar stress, a hypothesis supported by the nuclear accumulation of NAT10 upon actinomycin D (ActD) treatment, a known RNA Pol I inhibitor. ActD treatment also led to the formation of γH2AX foci, reinforcing the notion of a functional crosstalk between DDR and nucleolar stress pathways (Figure 2F–2G, S3F–S3G). Similar changes were detected for the nucleolar protein TCOF1, a validated DUSP12 interactor under genotoxic stress in tumor cells [46]. Confocal coefficient correlation obtained from microscopy analysis revealed enhanced colocalization of DUSP12 and NAT10 upon DX treatment in both cell lines (Figure 2I, S3I).

**Figure 2.**
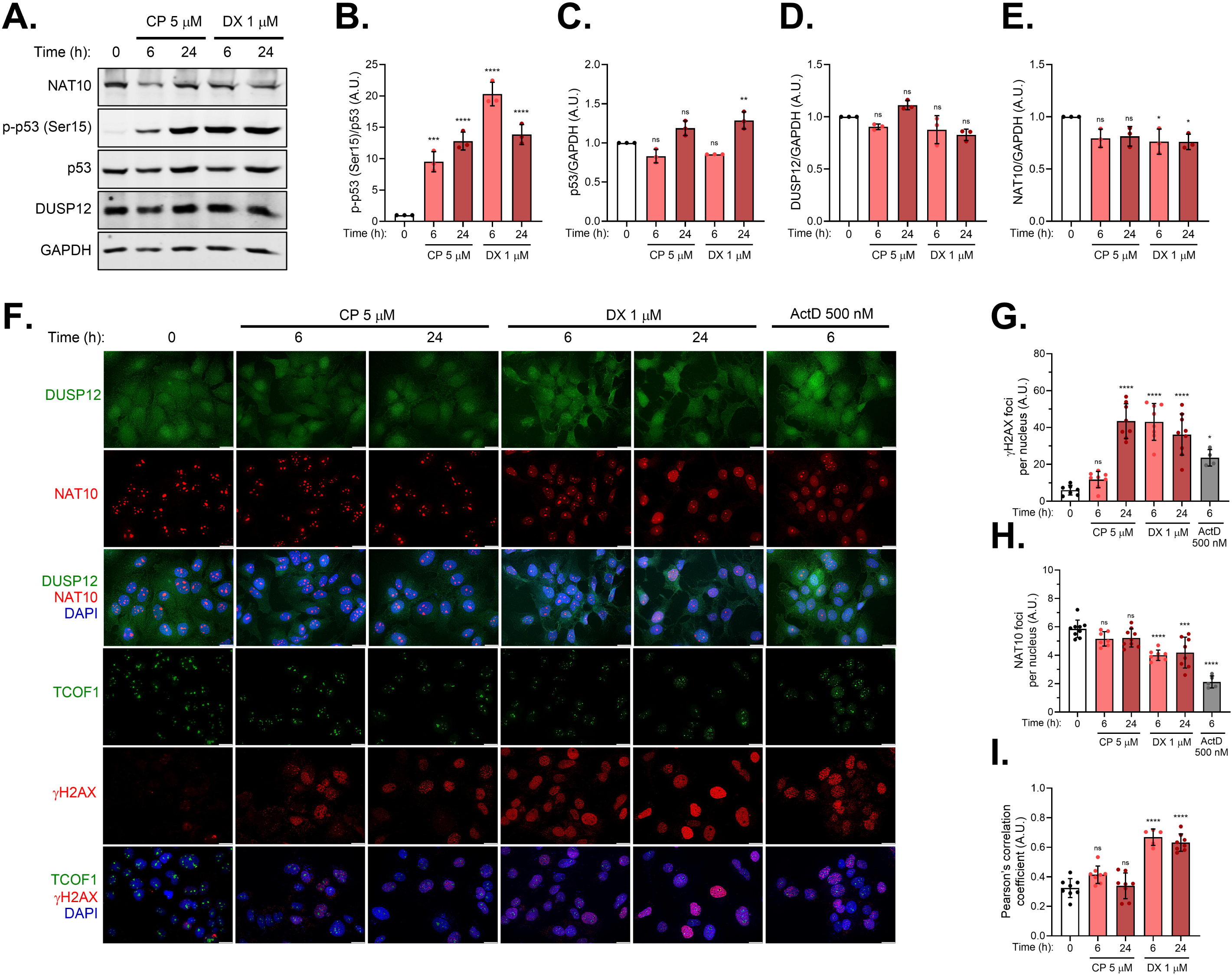
Doxorubicin triggers stronger DDR activation and NAT10 relocation in HuH-7 cells. (A) Western blot detection of NAT10, phospho-p53 (Ser15), total p53, DUSP12, and GAPDH in HuH-7 cells treated with CP or DX. (B–E) Densitometric quantification of (B) phospho-p53 (Ser15), (C) total p53, (D) DUSP12, and (E) NAT10 relative to GAPDH expression. (F) Representative immunofluorescence images of DUSP12, NAT10, TCOF1, and γH2AX in HuH-7 cells treated with CP, DX, or ActD. Scale bar = 20 μm. (G) Quantification of γH2AX foci per nucleus. (H) Quantification of NAT10 foci per nucleus. (I) Colocalization of DUSP12 and NAT10 assessed by Pearson’s correlation coefficient. Data represent mean ± SD of at least three independent experiments. Statistical significance relative to untreated cells is indicated as *p<0.05; **p<0.01; ***p<0.001; ****p<0.0001, ns = not significant.

### DUSP12 knockout sensitizes HCC cells and spheroids to doxorubicin

To further investigate the role of DUSP12 in HCC, CRISPR-Cas9 was employed to generate knockout (KO) clones with varying degrees of DUSP12 loss (Figure 3A–3B). No changes were observed in the ability of DUSP12-KO cells to form spheroids, although the resulting spheroids were consistently smaller in size (Figure S5A–S5D). Similarly, proliferation rates in both monolayer and spheroid cultures remained largely unaffected (Figure 3C, S6A). DUSP12-deficient cells exhibited impaired S-phase entry, as indicated by G1 accumulation (Figure 3D). In monolayer cultures, DUSP12 loss did not alter CP responsiveness, but markedly increased sensitivity to DX in a manner proportional to DUSP12 expression levels (Figure 3E–3F, Supplementary Table S7). Notably, DUSP12-KO spheroids were more sensitive to both CP and DX (Figure 3G–3H, S6B–S6C, Supplementary Table S8). Similarly, proliferation curves showed that while doubling time remained unchanged after CP treatment, DX-treated knockout cells displayed a trend of increasing doubling time with reduced DUSP12 expression (Figure 3I–3J, S6D–S6I, Supplementary Table S9).

**Figure 3.**
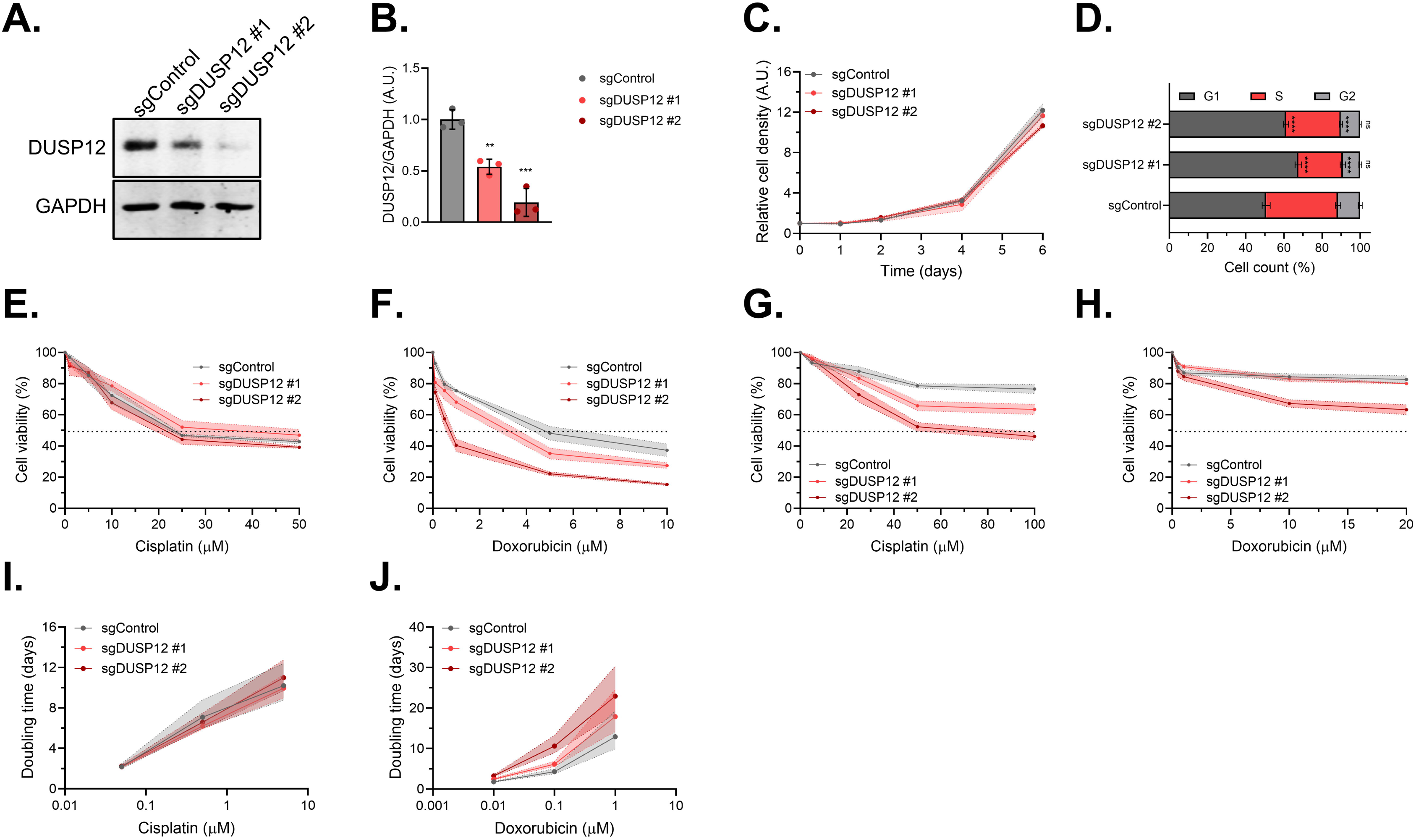
CRISPR-mediated knockout of DUSP12 sensitizes HuH-7 cells to doxorubicin. (A) Immunoblot validation of DUSP12 knockout in HuH-7 cells using two different sgRNAs (sgDUSP12 #1 and #2). (B) Densitometric quantification of DUSP12 relative to GAPDH expression. (C) Proliferation curves of sgControl and sgDUSP12 HuH-7 clones in monolayer cultures. (D) Cell cycle distribution of DUSP12-KO HuH-7 clones. Statistical significance relative to sgControl is indicated as *p<0.05; **p<0.01; ***p<0.001; ****p<0.0001, ns = not significant. (E–F) Cell viability of DUSP12-KO HuH-7 clones in monolayer cultures treated with increasing concentrations of (E) CP or (F) DX for 48 h. (G–H) Cell viability of DUSP12-KO HuH-7 spheroids treated with increasing concentrations of (G) CP or (H) DX for 48 or 72 h. (I–J) Doubling time of DUSP12-KO HuH-7 clones in monolayer cultures treated with increasing concentrations of (I) CP or (J) DX for 24 h. Graphs represent mean ± SD of at least three independent experiments.

### DUSP12 loss sensitizes HuH-7 cells to doxorubicin through impaired DNA repair, enhanced nucleolar stress, and synthetic lethality with NAT10 inhibition

To elucidate the mechanism underlying the increased sensitivity to doxorubicin observed in DUSP12 knockout cells, we first examined the two primary modes of action of doxorubicin: generation of reactive oxygen species (ROS) and induction of DNA damage. While no differences in ROS levels were detected across a wide range of doxorubicin concentrations between DUSP12-proficient and -deficient clones (Figure S7A), a notably stronger DDR activation was observed in the knockout clones, as evidenced by increased p53 expression and phosphorylation (Figure 4A–4C) and by the quantification of γH2AX foci (Figure 4D, S7B). The pronounced DDR activation observed following DX treatment appears to be a consequence of reduced repair capacity or delayed repair kinetics of both double-strand and single-strand DNA breaks in DUSP12-KO cells, as demonstrated by neutral and alkaline comet assays, respectively (Figure 4E–4F). Interestingly, DUSP12 knockout cells display features consistent with nucleolar stress, such as elevated basal levels of total p53 (Figure 4A, 4C) and fewer nucleoli per nucleus, as demonstrated by both NAT10 and TCOF1 staining (Figure 4G–4I). Upon DX treatment, DUSP12-proficient cells exhibited pronounced nucleolar remodeling characterized by NAT10 and TCOF1 redistribution from nucleoli to the nucleoplasm, leading to a marked reduction in nucleolar number and formation of nucleolar caps (Figure 4I). In contrast, DUSP12-deficient cells displayed altered nucleolar structure and dynamics: although NAT10 redistributed to the nucleoplasm, nucleolar number remained largely unchanged, and TCOF1 was strongly displaced to the nucleoplasm, losing its spatial association with NAT10-positive nucleoli (Figure 4G–4I). Given this altered nucleolar dynamics in DUSP12-deficient clones, we hypothesized that pharmacological inhibition of NAT10 could further enhance their sensitivity to genotoxic agents. Consistent with this hypothesis, inhibition of NAT10 with remodelin (RMD) did not sensitize cells to cisplatin but markedly increased sensitivity to doxorubicin (Figure 4J–4K). Although a modest effect was also observed in DUSP12-proficient cells, the reduction in IC_50_ values was considerably more pronounced in DUSP12 knockout clones (Supplementary Table S10). Moreover, all clones displayed comparable sensitivity to RMD alone (Figure S7C, Supplementary Table S11), suggesting that the combined inhibition of DUSP12 and NAT10 may result in a form of synthetic lethality selectively triggered by doxorubicin-induced genotoxic stress.

**Figure 4.**
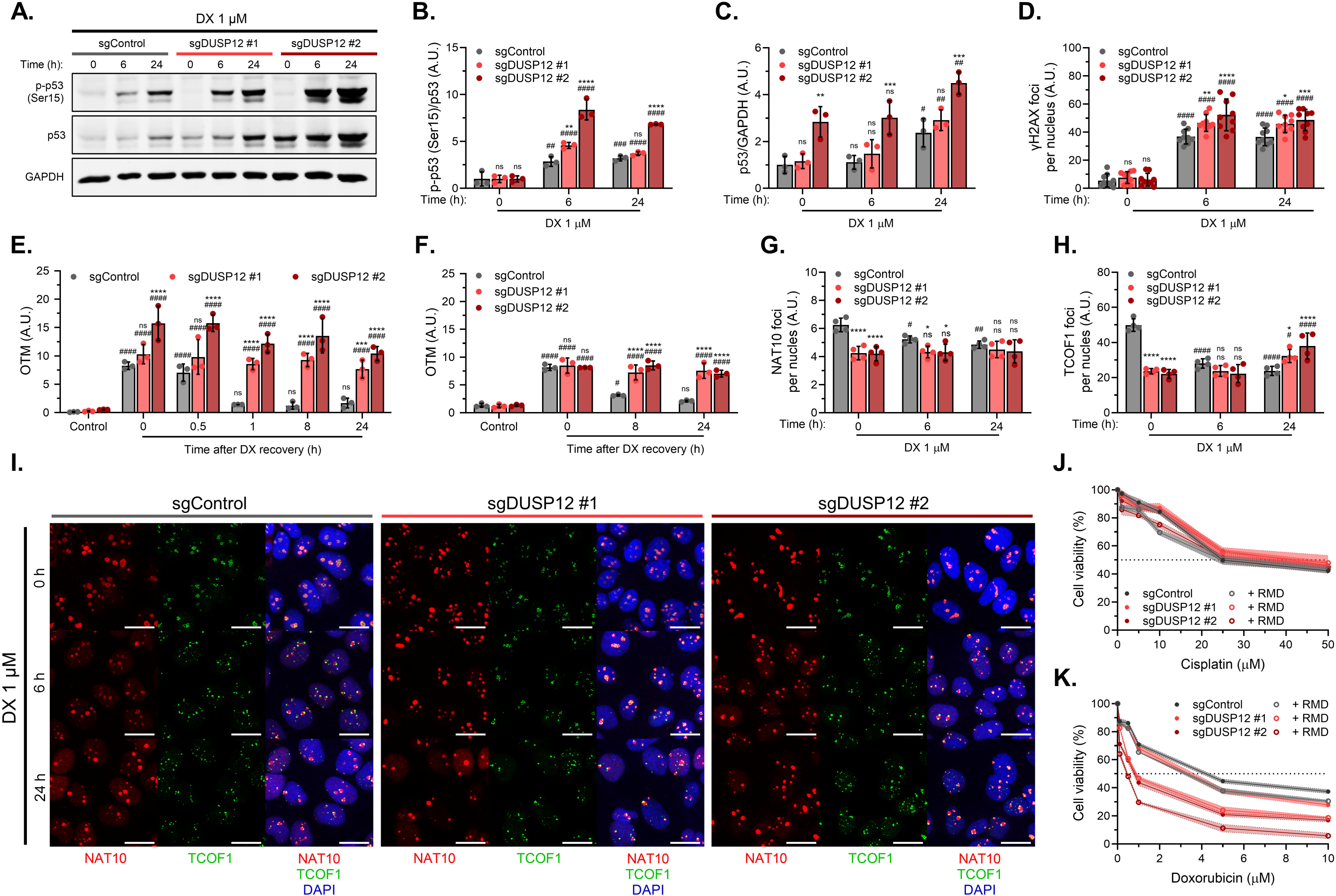
DUSP12 knockout clones exhibit enhanced DDR activation, impaired DNA repair, and enhanced nucleolar stress features after genotoxic stress. (A) Western blot detection of phospho-p53 (Ser15), total p53, and GAPDH in DUSP12-KO HuH-7 clones treated with DX under the indicated conditions. (B–C) Densitometric quantification of (B) phospho-p53 (Ser15) and (C) total p53 relative to GAPDH expression. (D) Quantification of γH2AX foci per nucleus. (E–F) DNA repair capacity assessed by Olive Tail Moment (OTM) using (E) alkaline and (F) neutral comet assays in DUSP12-KO HuH-7 clones after 24 h of DX treatment and subsequent recovery at the indicated times. (G) Quantification of NAT10 foci per nucleus. (H) Quantification of TCOF1 foci per nucleus. (I) Representative immunofluorescence images of NAT10 and TCOF1 in DUSP12-KO HuH-7 clones treated with DX. Scale bar = 5 μm. (J–K) Cell viability of DUSP12-KO HuH-7 clones pre-treated with 10 µM RMD for 24 h prior to exposure to increasing concentrations of (J) CP or (K) DX for 48 h (in the presence of RMD). Graphs represent mean ± SD of at least three independent experiments. Statistical significance relative to sgControl at each timepoint is indicated above as *p<0.05; **p<0.01; ***p<0.001; ****p<0.0001, ns = not significant. Statistical significance relative to untreated cells is indicated below as #p<0.05; ##p<0.01; ###p<0.001; ####p<0.0001, ns = not significant.

### Bidirectional regulation between DUSP12 and NAT10 modulates NAT10 phosphorylation and RNA acetylation

To gain insights into the interaction between DUSP12 and NAT10, pull-down assays were performed using recombinant GST-tagged DUSP12 constructs [46]. The full-length and the substrate-trapping mutant C115S displayed comparable binding to NAT10; in contrast, binding was significantly reduced when using the isolated N-terminal domain, showing that removal of the C-terminal zinc-binding domain compromises the interaction, although the N-terminal domain alone retains partial binding capacity (Figure 5A–5B). To assess whether DUSP12 catalytic activity contributes to complex formation, pull-down assays were performed in the presence of the tyrosine phosphatase inhibitor Na_3_VO_4_. After establishing the optimal inhibition conditions for DUSP12 inhibition by monitoring dephosphorylation of the general substrate DiFMUP (6,8-Difluoro-4-Methylumbelliferyl Phosphate) (Figure S8A), pull-down assay revealed that the interaction with both the full-length and N-terminal constructs was significantly reduced, indicating that DUSP12 catalytic activity is required for efficient interaction with NAT10, while inhibition had no effect on binding to the isolated C-terminal domain as expected (Figure S8B–S8C). Interestingly, treatment with RMD selectively reduced binding to full-length DUSP12 without affecting interaction with isolated domains (Figure S8D–S8E), suggesting that both domains – or their interface – are required for interaction with catalytically active NAT10. This supports a bidirectional mechanism in which both enzymes influence each other’s activity. Consistently, NAT10 immunoprecipitation followed by phosphotyrosine detection revealed increased NAT10 phosphorylation in sgDUSP12 #2 cells, supporting a direct role for DUSP12 in NAT10 dephosphorylation under basal conditions (Figure 5C–5D). Loss or reduction of DUSP12 also led to decreased levels of acetylated cytidine (ac4C) in RNA, suggesting impaired NAT10 catalytic activity (Figure 5E–5F). Finally, interaction between DUSP12 and NAT10 was reduced in spheroids and under genotoxic stress (CP or DX), suggesting stress-induced post-translational or conformational changes in NAT10 that impair complex formation (Figure 5G–5H).

**Figure 5.**
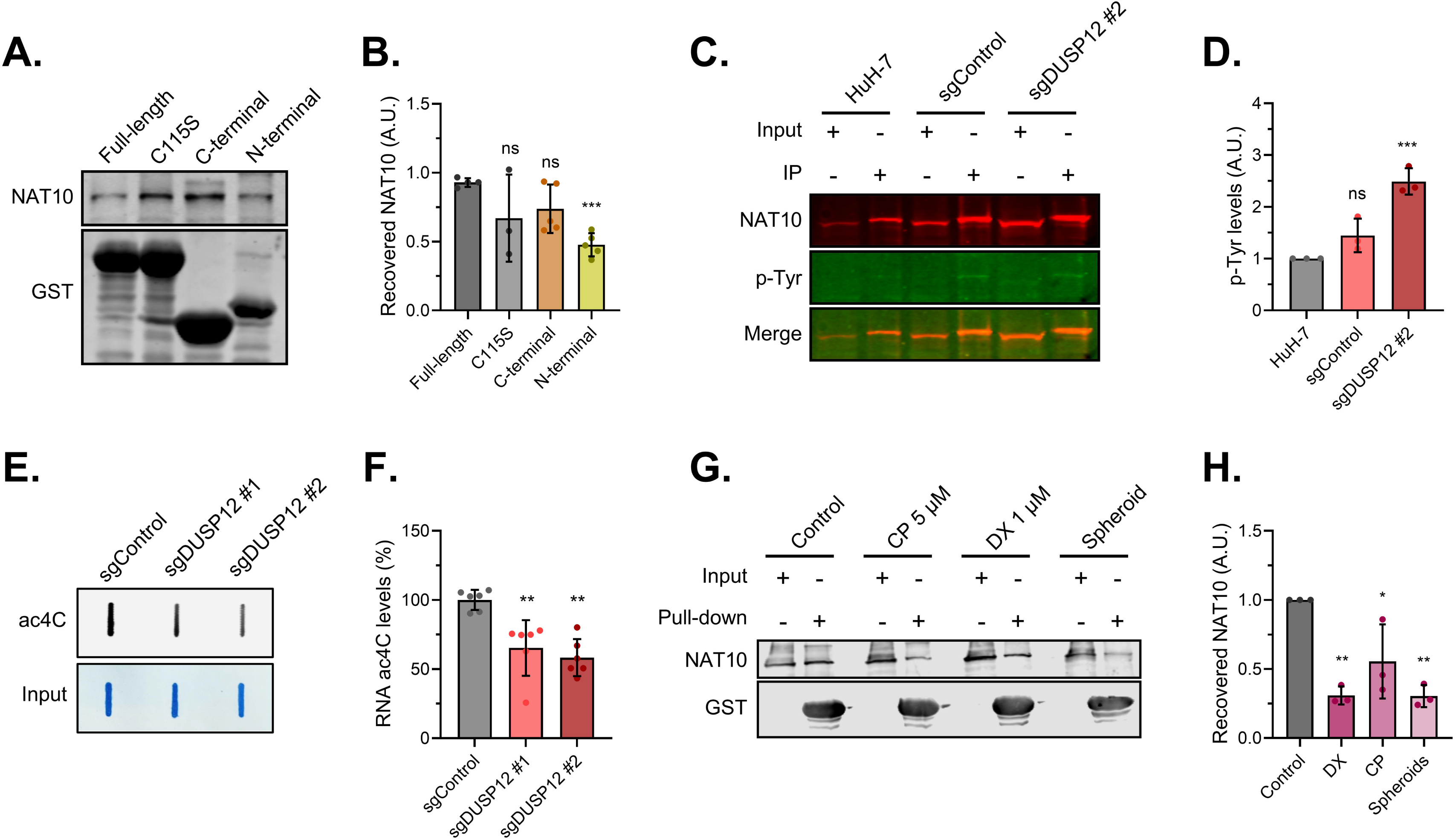
DUSP12 interacts with and dephosphorylates NAT10 to regulate its RNA acetyltransferase activity. (A) Western blot detection of NAT10 and GST after pull-down assays with GST-tagged DUSP12 constructs. (B) Densitometric quantification of NAT10 binding levels. Differences relative to full-length DUSP12 are indicated. (C) Western blot detection of NAT10 and phosphotyrosine (p-Tyr) after immunoprecipitation of NAT10 in HuH-7 parental cells, sgControl, and sgDUSP12 #2 clones. (D) Densitometric quantification of p-Tyr levels. Differences relative to parental HuH-7 cells are indicated. (E) Slot blot detection of ac4C in DUSP12-KO clones. (F) Densitometric quantification of ac4C levels. Differences relative to sgControl are indicated. G) Western blot detection of NAT10 from pull-down assays using full-length DUSP12 as bait incubated with lysates from HuH-7 cells in monolayer untreated or treated with 5 µM CP or 1 µM DX for 24 h, and from HuH-7 spheroid cultures. H) Densitometric quantification of NAT10 binding levels. Differences relative to HuH-7 cells in monolayer are indicated. Graphs represent mean ± SD of at least three independent experiments. Statistical significance is indicated as *p<0.05; **p<0.01; ***p<0.001; ****p<0.0001, ns = not significant.

### DUSP12 amplification correlates with poor prognosis and enrichment in ribosome biogenesis and DDR pathways in HCC patients

As a proof-of-concept about the roles of DUSP12 in HCC, patient-derived datasets were analyzed. The *dusp12* gene was frequently gained (10%) or amplified (63%) in HCC cases (Figure 6A). As expected, *dusp12* copy number gain was correlated with aneuploidy score (Figure S9A) and tended to increase with tumor grade (Figure S9B). Copy number alterations corresponded to elevated DUSP12 expression (Figure 6B), which was associated with reduced overall and disease-specific survival (Figure 6C–6D), but not with disease progression or recurrence (Figure S9C–S9D). Genes strongly co-expressed with DUSP12 (n = 1189, ρ > 0.3) were enriched in pathways related to both canonical functions of DUSP12, such as ribosome and ribonucleoprotein complex biogenesis, and non-canonical roles, including DNA replication initiation, kinetochore assembly, telomere maintenance, DNA repair, and chromatin remodeling (Figure 6E). Consistently, enrichment was also observed for cellular components such as the nucleoplasm, kinetochore, and heterochromatin, as well as molecular functions related to nucleic acid interactions (Figure S9E–S9F). Only 1.1% of co-expressed genes were located near the *dusp12* locus (cytogenetic band 1q23.3), suggesting that co-expression reflected functional rather than positional effects (Figure S9G). Notably, 71 out of the 400 nuclear interaction partners of DUSP12 identified under genotoxic stress in our laboratory [46] were also found among the co-expressed genes, including NAT10 and TCOF1 (Figure 6F–6G, S9H).

**Figure 6.**
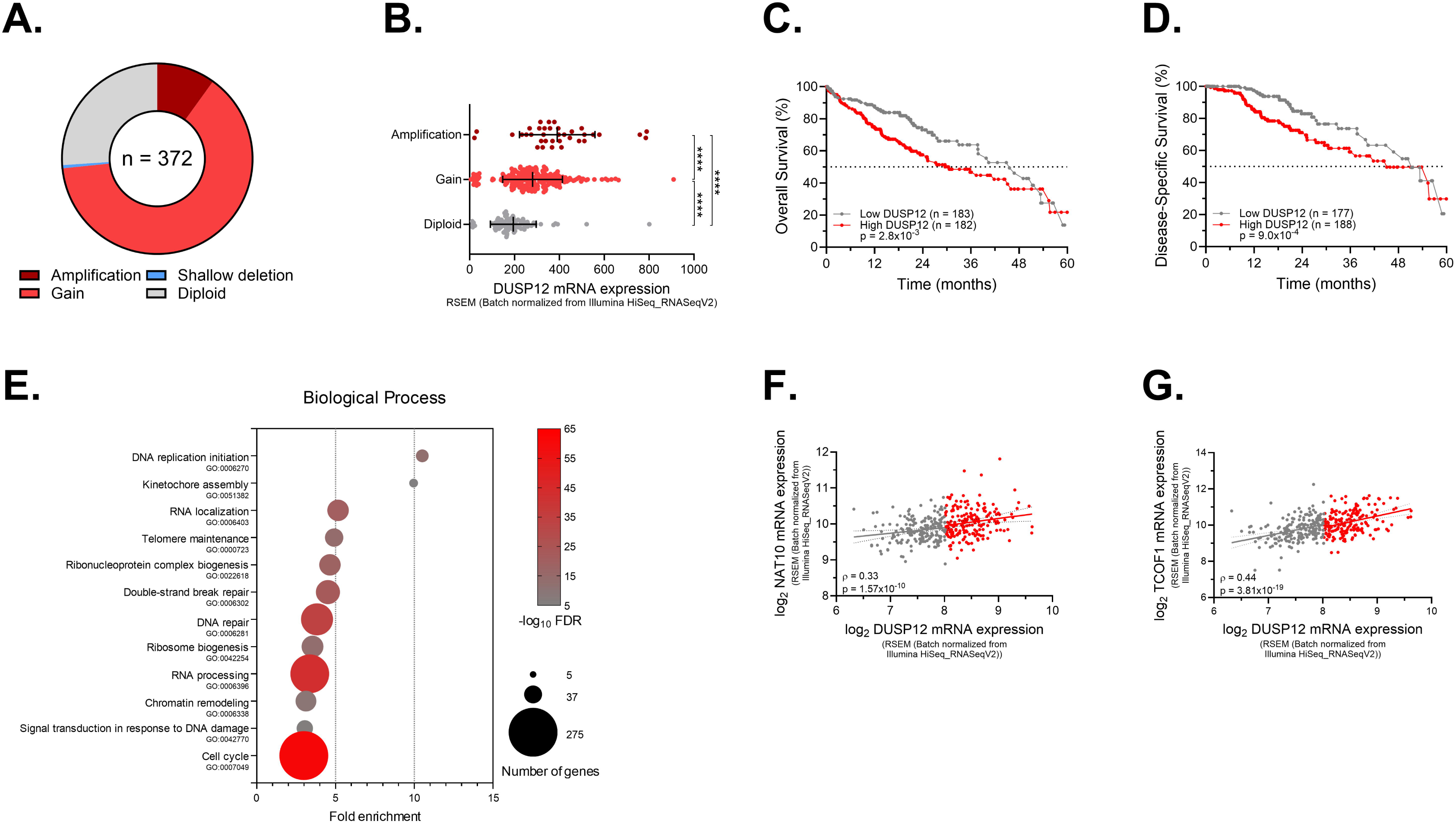
Genomic alterations and clinical significance of DUSP12 in HCC samples from human patients. (A) Distribution of *dusp12* copy number alterations in HCC patients. (B) DUSP12 mRNA expression in HCC patients with or without copy number alterations. (C–D) 5-year Kaplan-Meier analyses of (C) overall survival and (D) disease-specific survival in HCC patients stratified by median DUSP12 expression. (E) Gene ontology analysis of biological processes associated with genes correlated with DUSP12 expression in HCC patients. (F–G) Correlation between DUSP12 expression and (F) NAT10 or (G) TCOF1 expression in HCC patients with high (red) or low (gray) DUSP12 expression. Spearman’s correlation coefficient (ρ) and associated p-value are shown. Statistical significance is indicated as *p<0.05; **p<0.01; ***p<0.001; ****p<0.0001, ns = not significant.

## Discussion

This study identifies DUSP12 as a critical modulator of the cellular response to genotoxic and nucleolar stress in hepatocellular carcinoma (HCC). We demonstrate that doxorubicin (DX) induces a more robust and earlier DNA damage response (DDR) than cisplatin (CP), evidenced by increased p53 phosphorylation and γH2AX foci formation. While DUSP12 expression remained stable under genotoxic stress, NAT10 was redistributed from the nucleolus to the nucleoplasm. This relocation, alongside substantial nucleolar morphology alterations and parallel changes in TCOF1 localization, is consistent with nucleolar stress. Supporting this, actinomycin D (ActD), a canonical RNA Pol I inhibitor, also promoted nucleolar NAT10 redistribution and triggered γH2AX foci, confirming functional crosstalk between DDR and nucleolar stress pathways. DX-induced nucleolar stress may be particularly pronounced due to its preferential intercalation into GC-rich rDNA regions, impairing replication and repair [49]. Furthermore, DUSP12 localization differed between non-tumorigenic (HepaRG) and malignant (HuH-7) cells, suggesting its subcellular distribution may be altered during malignant transformation.

Loss-of-function experiments established a functional interplay between DUSP12 and chemoresistance. DUSP12 knockout sensitized HuH-7 cells to DX without affecting ROS levels, indicating a DDR-related mechanism. This complements previous models of DUSP12 in oxidative stress survival [27,45,51] by demonstrating its role in modulating genotoxic stress responses. Despite residual expression—potentially due to the high aneuploidy of HuH-7 cells and frequent DUSP12 copy number gains in HCC—DUSP12-deficient cells exhibited enhanced DDR activation. This suggests delayed or inefficient repair of DNA breaks [52]. The marked G1 arrest in knockout cells indicates reduced opportunity for homologous recombination (HR), potentially shifting repair toward error-prone non-homologous end joining (NHEJ), which could partly explain the increased DX sensitivity.

Beyond DDR modulation, DUSP12 loss elicited features of nucleolar stress, including increased basal p53 levels and altered nucleolar organization. This aligns with DUSP12’s role in ribosome biogenesis and its physical interaction with nucleolar regulators [18,20–23,46]. Notably, DUSP12 knockout disrupted the localization of its validated partners, NAT10 and TCOF1, redistributing them to the nucleoplasm even without genotoxic challenge, with further mislocalization following DX treatment. This is significant given that both proteins are upregulated in HCC and linked to therapeutic resistance [37,53]. TCOF1 acts as a central coordinator of nucleolar DDR, recruiting the ATR activator TOPBP1 to nucleoli following rDNA breaks in an ATM/NBS1-dependent manner, which is essential for inhibiting rRNA synthesis and nucleolar remodeling [54–56]. The pronounced TCOF1 redistribution in DUSP12-deficient cells likely reflects impaired DDR coordination within the nucleolus and defective stress adaptation. Therapeutically, this state is exploitable, as DUSP12 loss markedly sensitized cells to DX, a drug that itself induces nucleolar remodeling, indicating that targeting nucleolar homeostasis could enhance chemotherapeutic efficacy in HCC. Critically, we uncovered a novel molecular connection between DUSP12 and NAT10. Immunoprecipitation confirmed physical interaction, and experimental evidence supports NAT10 as a potential substrate for DUSP12-mediated dephosphorylation. Inhibition of either phosphatase (Na3VO4) or acetyltransferase (Remodelin, RMD) activity reduced this interaction, suggesting a bidirectional regulatory mechanism. Consistently, DUSP12 loss increased NAT10 phosphotyrosine levels and impaired RNA acetylation (ac4C). While ac4C is essential for 18S rRNA maturation in some organisms, it appears dispensable in human cells, potentially explaining why DUSP12 loss alone does not compromise proliferation [57]. However, ac4C is implicated in DNA repair: it stabilizes AHNAK mRNA to recruit LIG4/XRCC4 during NHEJ, destabilizes DDB2 mRNA to attenuate global genome nucleotide excision repair (GG-NER) [58,59], and NAT10 accumulation at double-strand breaks promotes HR via ac4C modification of RNAs in DNA:RNA hybrids, likely through chromatin decondensation [60,61]. In line with this, pharmacological inhibition of NAT10 with RMD synergistically enhanced DX sensitivity in DUSP12-KO cells, indicating a synthetic-lethal interaction specific to this axis under DX-induced stress. The lack of synergy with cisplatin underscores a DNA lesion-specific mechanism.

Finally, analysis of patient-derived datasets confirmed the clinical relevance of our findings. DUSP12 is frequently amplified in HCC, correlates with NAT10 expression and poor prognosis, and is co-expressed with genes enriched in ribosome biogenesis, DDR, and chromatin-related pathways. Recent studies show NAT10 promotes HCC progression by stabilizing oncogenic transcripts like DDIAS via ac4C [62], and ac4C-related mRNA signatures stratify HCC patients into prognostic groups, with higher ac4C associating with aggressive features, increased DNA repair activation, and poorer outcomes, including differential responses to immunotherapy and mTOR inhibition [63].y These observations reinforce the physiological relevance of ac4C in DDR and therapeutic sensitivity.

In conclusion, our data position DUSP12 upstream of NAT10, linking tyrosine dephosphorylation to RNA acetylation control. The DUSP12-NAT10-ac4C axis is a critical determinant of genome stability, nucleolar function, and therapeutic response, establishing DUSP12 as a promising candidate for targeted intervention in HCC.

## Acknowledgments

The authors thank Prof. Alexandre B. Cardoso (IQ-USP) for allowing the use of the Leica DMi1 and DMi8 microscopes, Prof. Pio C. Neto and Prof. Ohara Augusto for allowing the use of Tecan equipment.

## Data availability statement

The results obtained in this study are presented in the main and supplementary figures. Other data not shown can be obtained by consulting the authors.

## Contributions

Conception and design: VKB, FLF. Development of methodology: VKB, FLF. Acquisition of data: VKB, YTM, DRDCGP, FLF. Analysis and interpretation of data: VKB, YTM, DRDCGP, IYAO, NCH, FLF. Writing, review, and/or revision of the manuscript: VKB, YTM, DRDCGP, IYAO, NCH, FLF. Administrative, technical, or material support: VKB, FLF. Fundings acquisition: FLF. Study supervision: FLF.

## References

[1] Vogel A, Meyer T, Sapisochin G, Salem R, Saborowski A. Hepatocellular carcinoma. The Lancet 2022;400:1345–62. 10.1016/S0140-6736(22)01200-4.

[2] Gordan JD, Kennedy EB, Ghassan;, Abou-Alfa K, Muhammad;, Beg S, et al. Systemic Therapy for Advanced Hepatocellular Carcinoma: ASCO Guideline. J Clin Oncol 2020;38:4317–45. 10.1200/JCO.20.

[3] Raoul J-L, Forner A, Bolondi L, Cheung TT, Kloeckner R, de Baere T. Updated use of TACE for hepatocellular carcinoma treatment: How and when to use it based on clinical evidence. Cancer Treat Rev 2019;72:28–36. 10.1016/j.ctrv.2018.11.002.

[4] Pelletier J, Thomas G, Volarević S. Ribosome biogenesis in cancer: new players and therapeutic avenues. Nat Rev Cancer 2018;18:51–63.

[5] Yang K, Wang M, Zhao Y, Sun X, Yang Y, Li X, et al. A redox mechanism underlying nucleolar stress sensing by nucleophosmin. Nat Commun 2016;7. 10.1038/ncomms13599.

[6] Chen H, Han L, Tsai H, Wang Z, Wu Y, Duo Y, et al. PICT-1 is a key nucleolar sensor in DNA damage response signaling that regulates apoptosis through the RPL11-MDM2-p53 pathway. vol. 7. 2016.

[7] Dai MS, Lu H. Inhibition of MDM2-mediated p53 ubiquitination and degradation by ribosomal protein L5. Journal of Biological Chemistry 2004;279:44475–82. 10.1074/jbc.M403722200.

[8] Lohrum MAE, Ludwig RL, Kubbutat MHG, Hanlon M, Vousden KH. Regulation of HDM2 activity by the ribosomal protein L11. Cancer Cell 2003;3:577–87.

[9] Sokka M, Rilla K, Miinalainen I, Pospiech H, Syväoja JE. High levels of TopBP1 induce ATR-dependent shut-down of rRNA transcription and nucleolar segregation. Nucleic Acids Res 2015;43:4975–89. 10.1093/nar/gkv371.

[10] Larsen DH, Hari F, Clapperton JA, Gwerder M, Gutsche K, Altmeyer M, et al. The NBS1-Treacle complex controls ribosomal RNA transcription in response to DNA damage. Nat Cell Biol 2014;16:792–803. 10.1038/ncb3007.

[11] Calkins AS, Iglehart JD, Lazaro JB. DNA damage-induced inhibition of rRNA synthesis by DNA-PK and PARP-1. Nucleic Acids Res 2013;41:7378–86. 10.1093/nar/gkt502.

[12] Kruhlak M, Crouch EE, Orlov M, Montão C, Gorski SA, Nussenzweig A, et al. The ATM repair pathway inhibits RNA polymerase I transcription in response to chromosome breaks. Nature 2007;447:730–4. 10.1038/nature05842.

[13] Azman MS, Alard EL, Dodel M, Capraro F, Faraway R, Dermit M, et al. An ERK1/2-driven RNA-binding switch in nucleolin drives ribosome biogenesis and pancreatic tumorigenesis downstream of RAS oncogene. EMBO J 2023;42. 10.15252/embj.2022110902.

[14] Iadevaia V, Zhang Z, Jan E, Proud CG. MTOR signaling regulates the processing of pre-rRNA in human cells. Nucleic Acids Res 2012;40:2527–39. 10.1093/nar/gkr1040.

[15] Cornelison R, Dobbin ZC, Katre AA, Jeong DH, Zhang Y, Chen D, et al. Targeting RNA-polymerase I in both chemosensitive and chemoresistant populations in epithelial ovarian cancer. Clinical Cancer Research 2017;23:6529–40. 10.1158/1078-0432.CCR-17-0282.

[16] Lawrence MG, Porter LH, Choo N, Pook D, Grummet JP, Pezaro CJ, et al. CX-5461 sensitizes DNA damage repair-proficient castrate-resistant prostate cancer to PARP inhibition. Mol Cancer Ther 2021;20:2140–50. 10.1158/1535-7163.MCT-20-0932.

[17] Ban Y, Zou Y, Liu Y, Lee S, Bednarczyk RB, Sheng J, et al. Targeting ribosome biogenesis as a novel therapeutic approach to overcome EMT-related chemoresistance in breast cancer. Elife 2024;12. 10.7554/eLife.89486.

[18] Geng Q, Xhabija B, Knuckle C, Bonham CA, Vacratsis PO. The Atypical Dual Specificity Phosphatase hYVH1 Associates with Multiple Ribonucleoprotein Particles. Journal of Biological Chemistry 2017;292:539–50. 10.1074/jbc.M116.715607.

[19] Muda M, Manning ER, Orth K, Dixon JE. Identification of the Human YVH1 Protein-tyrosine Phosphatase Orthologue Reveals a Novel Zinc Binding Domain Essential for in Vivo Function*. J Biol Chem 1999;274:23991–5.

[20] Lo KY, Li Z, Wang F, Marcotte EM, Johnson AW. Ribosome stalk assembly requires the dual-specificity phosphatase Yvh1 for the exchange of Mrt4 with P0. Journal of Cell Biology 2009;186:849–62. 10.1083/jcb.200904110.

[21] Liang X, Zuo MQ, Zhang Y, Li N, Ma C, Dong MQ, et al. Structural snapshots of human pre-60S ribosomal particles before and after nuclear export. Nat Commun 2020;11. 10.1038/s41467-020-17237-x.

[22] Lo KY, Li Z, Bussiere C, Bresson S, Marcotte EM, Johnson AW. Defining the pathway of cytoplasmic maturation of the 60S ribosomal subunit. Mol Cell 2010;39:196–208. 10.1016/j.molcel.2010.06.018.

[23] DaDalt AA, Bonham CA, Lotze GP, Luiso AA, Vacratsis PO. Src-mediated phosphorylation of the ribosome biogenesis factor hYVH1 affects its localization, promoting partitioning to the 60S ribosomal subunit. Journal of Biological Chemistry 2022;298. 10.1016/j.jbc.2022.102679.

[24] Li H, Yang Q, Huang Z, Liang C, Zhang DH, Shi HT, et al. Dual-specificity phosphatase 12 attenuates oxidative stress injury and apoptosis in diabetic cardiomyopathy via the ASK1-JNK/p38 signaling pathway. Free Radic Biol Med 2022;192:13–24. 10.1016/j.freeradbiomed.2022.09.004.

[25] He J, Li S, Teng Y, Xiong H, Wang Z, Han X, et al. Increasing expression of dual-specificity phosphatase 12 mitigates oxygen-glucose deprivation/reoxygenation-induced neuronal apoptosis and inflammation through inactivation of the ASK1-JNK/p38 MAPK pathway. Autoimmunity 2024;57. 10.1080/08916934.2024.2345919.

[26] Huang Z, Wu L-M, Zhang J-L, Sabri A, Wang S-J, Qin G-J, et al. Dual Specificity Phosphatase 12 Regulates Hepatic Lipid Metabolism Through Inhibition of the Lipogenesis and Apoptosis Signal–Regulating Kinase 1 Pathways. Hepatology 2019;70:1099–118. 10.1002/hep.30597/suppinfo.

[27] Qiu T, Wang T, Zhou J, Chen Z, Zou J, Zhang L, et al. DUSP12 protects against hepatic ischemia–reperfusion injury dependent on ASK1-JNK/p38 pathway in vitro and in vivo. Clin Sci 2020;134:2279–94. 10.1042/CS20191272.

[28] Monteiro LF, Forti FL. Network analysis of DUSP12 partners in the nucleus under genotoxic stress. J Proteomics 2019;197:42–52. 10.1016/j.jprot.2019.02.008.

[29] Arango D, Sturgill D, Alhusaini N, Dillman AA, Sweet TJ, Hanson G, et al. Acetylation of Cytidine in mRNA Promotes Translation Efficiency. Cell 2018;175:1872–1886.e24. 10.1016/j.cell.2018.10.030.

[30] Sharma S, Langhendries JL, Watzinger P, Kotter P, Entian KD, Lafontaine DLJ. Yeast Kre33 and human NAT10 are conserved 18S rRNA cytosine acetyltransferases that modify tRNAs assisted by the adaptor Tan1/THUMPD1. Nucleic Acids Res 2015;43:2242–58. 10.1093/nar/gkv075.

[31] Suzuki T, Ito S, Horikawa S, Suzuki T, Kawauchi H, Tanaka Y, et al. Human NAT10 is an ATP-dependent rna acetyltransferase responsible for N4-acetylcytidine formation in 18 S ribosomal RNA (rRNA). Journal of Biological Chemistry 2014;289:35724–30. 10.1074/jbc.C114.602698.

[32] Cai S, Liu X, Zhang C, Xing B, Du X. Autoacetylation of NAT10 is critical for its function in rRNA transcription activation. Biochem Biophys Res Commun 2017;483:624–9. 10.1016/j.bbrc.2016.12.092.

[33] Hu Z, Lu Y, Cao J, Lin L, Chen X, Zhou Z, et al. N-acetyltransferase NAT10 controls cell fates via connecting mRNA cytidine acetylation to chromatin signaling. Sci Adv 2024;10:9871.

[34] Liu HY, Liu YY, Zhang YL, Ning Y, Zhang FL, Li DQ. Poly(ADP-ribosyl)ation of acetyltransferase NAT10 by PARP1 is required for its nucleoplasmic translocation and function in response to DNA damage. Cell Communication and Signaling 2022;20. 10.1186/s12964-022-00932-1.

[35] Liu X, Cai S, Zhang C, Liu Z, Luo J, Xing B, et al. Deacetylation of NAT10 by Sirt1 promotes the transition from rRNA biogenesis to autophagy upon energy stress. Nucleic Acids Res 2018;46:9601–16. 10.1093/nar/gky777.

[36] Zheng J, Tan Y, Liu X, Zhang C, Su K, Jiang Y, et al. NAT10 regulates mitotic cell fate by acetylating Eg5 to control bipolar spindle assembly and chromosome segregation. Cell Death Differ 2022;29:846–60. 10.1038/s41418-021-00899-5.

[37] Wang Y, Su K, Wang C, Deng T, Liu X, Sun S, et al. Chemotherapy-induced acetylation of ACLY by NAT10 promotes its nuclear accumulation and acetyl-CoA production to drive chemoresistance in hepatocellular carcinoma. Cell Death Dis 2024;15. 10.1038/s41419-024-06951-9.

[38] Liu H, Xu L, Yue S, Su H, Chen X, Liu Q, et al. Targeting N4-acetylcytidine suppresses hepatocellular carcinoma progression by repressing eEF2-mediated HMGB2 mRNA translation. Cancer Commun 2024. 10.1002/cac2.12595.

[39] Tan Y, Zheng J, Liu X, Lu M, Zhang C, Xing B, et al. Loss of nucleolar localization of NAT10 promotes cell migration and invasion in hepatocellular carcinoma. Biochem Biophys Res Commun 2018;499:1032–8. 10.1016/j.bbrc.2018.04.047.

[40] Friedrich J, Seidel C, Ebner R, Kunz-Schughart LA. Spheroid-based drug screen: considerations and practical approach. Nat Protoc 2009;4:309–24. 10.1038/nprot.2008.226.

[41] Friedrich J, Eder W, Castaneda J, Doss M, Huber E, Ebner R, et al. A Reliable Tool to Determine Cell Viability in Complex 3-D Culture: The Acid Phosphatase Assay. SLAS Discovery 2007;12:925–37. 10.1177/1087057107306839.

[42] Magalhaes YT, Forti FL. ROCK inhibition reduces the sensitivity of mutant p53 glioblastoma to genotoxic stress through a Rac1-driven ROS production. Int J Biochem Cell Biol 2023;164:106474. 10.1016/j.biocel.2023.106474.

[43] Mambetsariev N, Lin WW, Stunz LL, Hanson BM, Hildebrand JM, Bishop GA. Nuclear TRAF3 is a negative regulator of CREB in B cells. Proceedings of the National Academy of Sciences 2016;113:1032–7. 10.1073/pnas.1514586113.

[44] Torres TEP, Russo LC, Santos A, Marques GR, Magalhaes YT, Tabassum S, et al. Loss of DUSP3 activity radiosensitizes human tumor cell lines via attenuation of DNA repair pathways. Biochimica et Biophysica Acta (BBA) - General Subjects 2017;1861:1879–94. 10.1016/j.bbagen.2017.04.004.

[45] Bonham CA, Vacratsis PO. Redox Regulation of the Human Dual Specificity Phosphatase YVH1 through Disulfide Bond Formation. Journal of Biological Chemistry 2009;284:22853–64. 10.1074/jbc.M109.038612.

[46] Monteiro LF, Forti FL. Network analysis of DUSP12 partners in the nucleus under genotoxic stress. J Proteomics 2019;197:42–52. 10.1016/j.jprot.2019.02.008.

[47] Gao J, Aksoy BA, Dogrusoz U, Dresdner G, Gross B, Sumer SO, et al. Integrative Analysis of Complex Cancer Genomics and Clinical Profiles Using the cBioPortal. Sci Signal 2013;6. 10.1126/scisignal.2004088.

[48] Ge SX, Jung D, Yao R. ShinyGO: a graphical gene-set enrichment tool for animals and plants. Bioinformatics 2020;36:2628–9. 10.1093/bioinformatics/btz931.

[49] Yang F, Teves SS, Kemp CJ, Henikoff S. Doxorubicin, DNA torsion, and chromatin dynamics. Biochimica et Biophysica Acta (BBA) - Reviews on Cancer 2014;1845:84–9. 10.1016/j.bbcan.2013.12.002.

[50] Ju G, Zhou T, Zhang R, Pan X, Xue B, Miao S. DUSP12 regulates the tumorigenesis and prognosis of hepatocellular carcinoma. PeerJ 2021;9. 10.7717/peerj.11929.

[51] Sharda PR, Bonham CA, Mucaki EJ, Butt Z, Vacratsis PO. The dual-specificity phosphatase hYVH1 interacts with Hsp70 and prevents heat-shock-induced cell death. Biochemical Journal 2009;418:391–401. 10.1042/BJ20081484.

[52] Kasai F, Hirayama N, Ozawa M, Satoh M, Kohara A. HuH-7 reference genome profile: complex karyotype composed of massive loss of heterozygosity. Hum Cell 2018;31:261–7. 10.1007/s13577-018-0212-3.

[53] Wu C, Xia D, Wang D, Wang S, Sun Z, Xu B, et al. TCOF1 coordinates oncogenic activation and rRNA production and promotes tumorigenesis in HCC. Cancer Sci 2022;113:553–64. 10.1111/cas.15242.

[54] Larsen DH, Hari F, Clapperton JA, Gwerder M, Gutsche K, Altmeyer M, et al. The NBS1-Treacle complex controls ribosomal RNA transcription in response to DNA damage. Nat Cell Biol 2014;16:792–803. 10.1038/ncb3007.

[55] Mooser C, Symeonidou IE, Leimbacher PA, Ribeiro A, Shorrocks AMK, Jungmichel S, et al. Treacle controls the nucleolar response to rDNA breaks via TOPBP1 recruitment and ATR activation. Nat Commun 2020;11. 10.1038/s41467-019-13981-x.

[56] Ciccia A, Huang J-W, Izhar L, Sowa ME, Harper JW, Elledge SJ. Treacher Collins syndrome TCOF1 protein cooperates with NBS1 in the DNA damage response. Proceedings of the National Academy of Sciences 2014;111:18631–6. 10.1073/pnas.1422488112.

[57] Bortolin-Cavaillé M-L, Quillien A, Thalalla Gamage S, Thomas JM, Sas-Chen A, Sharma S, et al. Probing small ribosomal subunit RNA helix 45 acetylation across eukaryotic evolution. Nucleic Acids Res 2022;50:6284–99. 10.1093/nar/gkac404.

[58] Xie R, Cheng L, Huang M, Huang L, Chen Z, Zhang Q, et al. NAT10 Drives Cisplatin Chemoresistance by Enhancing ac4C-Associated DNA Repair in Bladder Cancer. Cancer Res 2023;83:1666–83. 10.1158/0008-5472.CAN-22-2233.

[59] Yang Z, Wilkinson E, Cui Y-H, Li H, He Y-Y. NAT10 regulates the repair of UVB-induced DNA damage and tumorigenicity. Toxicol Appl Pharmacol 2023;477:116688. 10.1016/j.taap.2023.116688.

[60] Svobodová Kovaříková A, Stixová L, Kovařík A, Bártová E. PARP-dependent and NAT10-independent acetylation of N4-cytidine in RNA appears in UV-damaged chromatin. Epigenetics Chromatin 2023;16:26. 10.1186/s13072-023-00501-x.

[61] Xu Z, Zhu M, Geng L, Zhang J, Xia J, Wang Q, et al. Targeting NAT10 attenuates homologous recombination via destabilizing DNA:RNA hybrids and overcomes PARP inhibitor resistance in cancers. Drug Resistance Updates 2025;81:101241. 10.1016/j.drup.2025.101241.

[62] Tao Y, Wang L, Chen E, Zhang S, Yang D, Chen W, et al. NAT10 promotes hepatocellular carcinoma progression by modulating the ac4C-DDIAS-PI3K-Akt axis. Sci Rep 2025;15:17286. 10.1038/s41598-025-00707-x.

[63] Liu S, Zhang Y, Qiu L, Zhang S, Meng Y, Huang C, et al. Uncovering N4-Acetylcytidine-Related mRNA Modification Pattern and Landscape of Stemness and Immunity in Hepatocellular Carcinoma. Front Cell Dev Biol 2022;10. 10.3389/fcell.2022.861000.

