## Supplementary Figures and Tables for "DUSP12 regulates NAT10-mediated RNA acetylation to modulate DNA repair and therapeutic response in hepatocellular carcinoma"

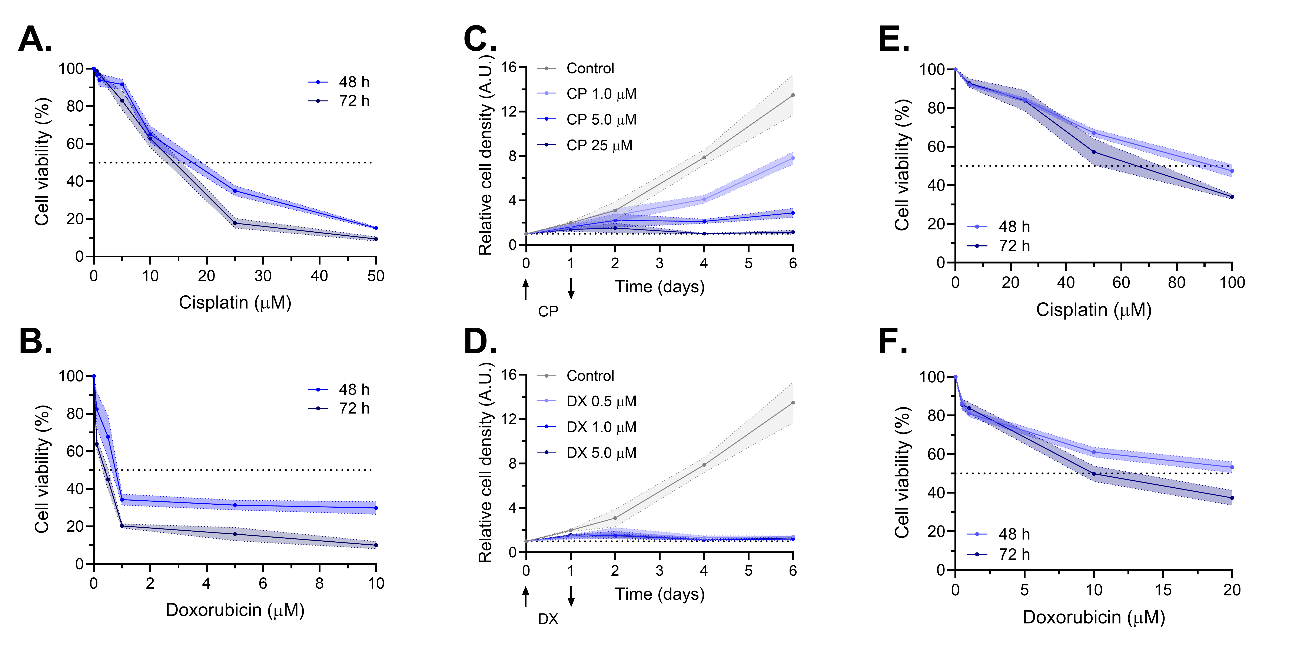

**Figure S1. Cytotoxic effects of cisplatin and doxorubicin in monolayers and spheroids of HepaRG cells.** (A–B) Cell viability of HepaRG monolayers treated with increasing concentrations of (A) CP or (B) DX for 48 or 72 h. (C–D) Proliferation curves of HepaRG monolayers treated with (C) CP or (D) DX for 24 h. (E–F) Cell viability of HepaRG spheroids treated with increasing concentrations of (E) CP or (F) DX for 48 or 72 h. Data represent mean ± SD of at least three independent experiments.

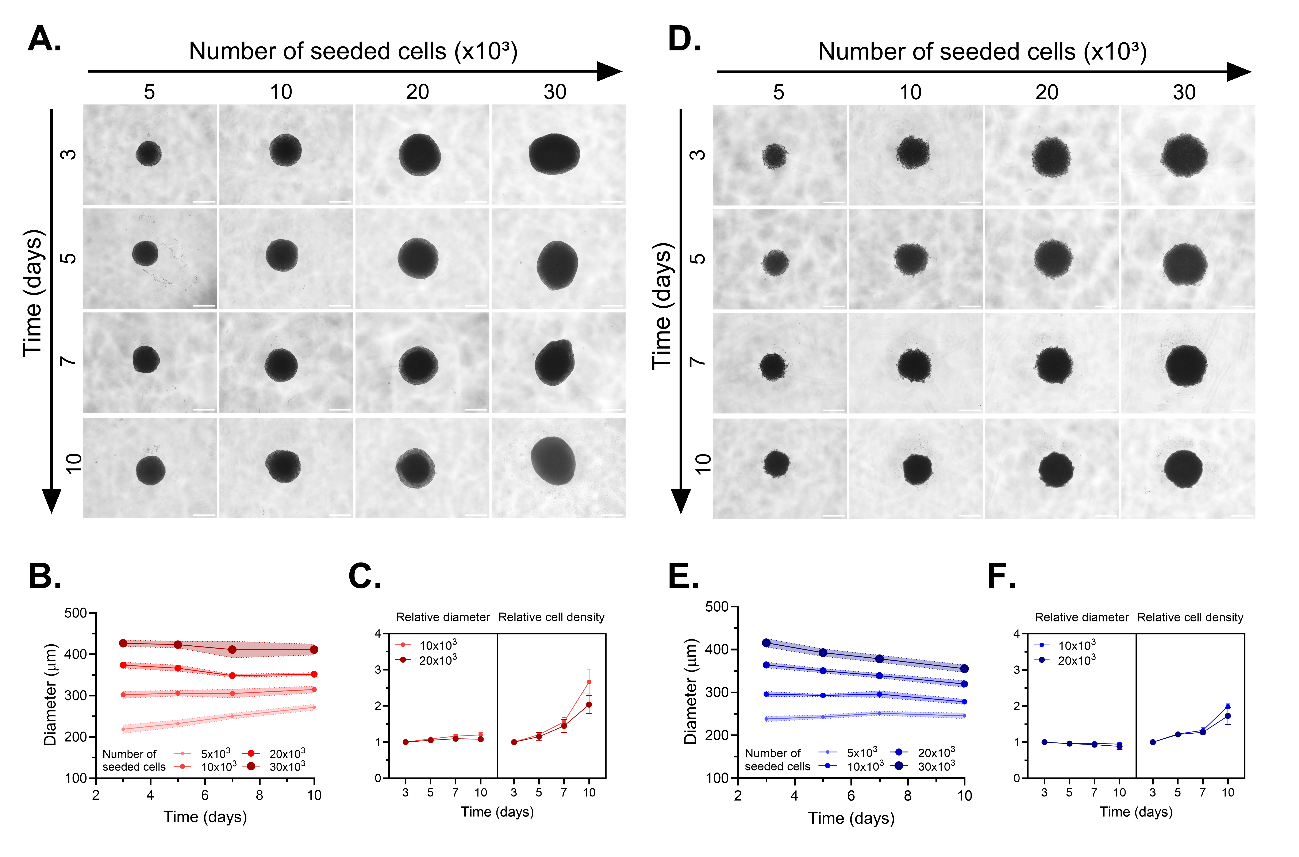

**Figure S2. Characterization of HuH-7 and HepaRG spheroid formation.** (A) Representative micrographs of HuH-7 spheroids generated by plating different cell numbers and maintained in culture for the indicated times. Scale bar = 200 μm. (B) Quantification of mean spheroid diameter (μm) under the conditions described above. (C) Relative variation in spheroid diameter and viability compared to day 3. (D) Representative micrographs of HepaRG spheroids generated by plating different cell numbers and maintained in culture for the indicated times. Scale bar = 200 μm. (E) Quantification of mean spheroid diameter (μm) under the conditions described above. (F) Relative variation in spheroid diameter and viability compared to day 3. Data represent mean ± SD of at least three independent experiments.

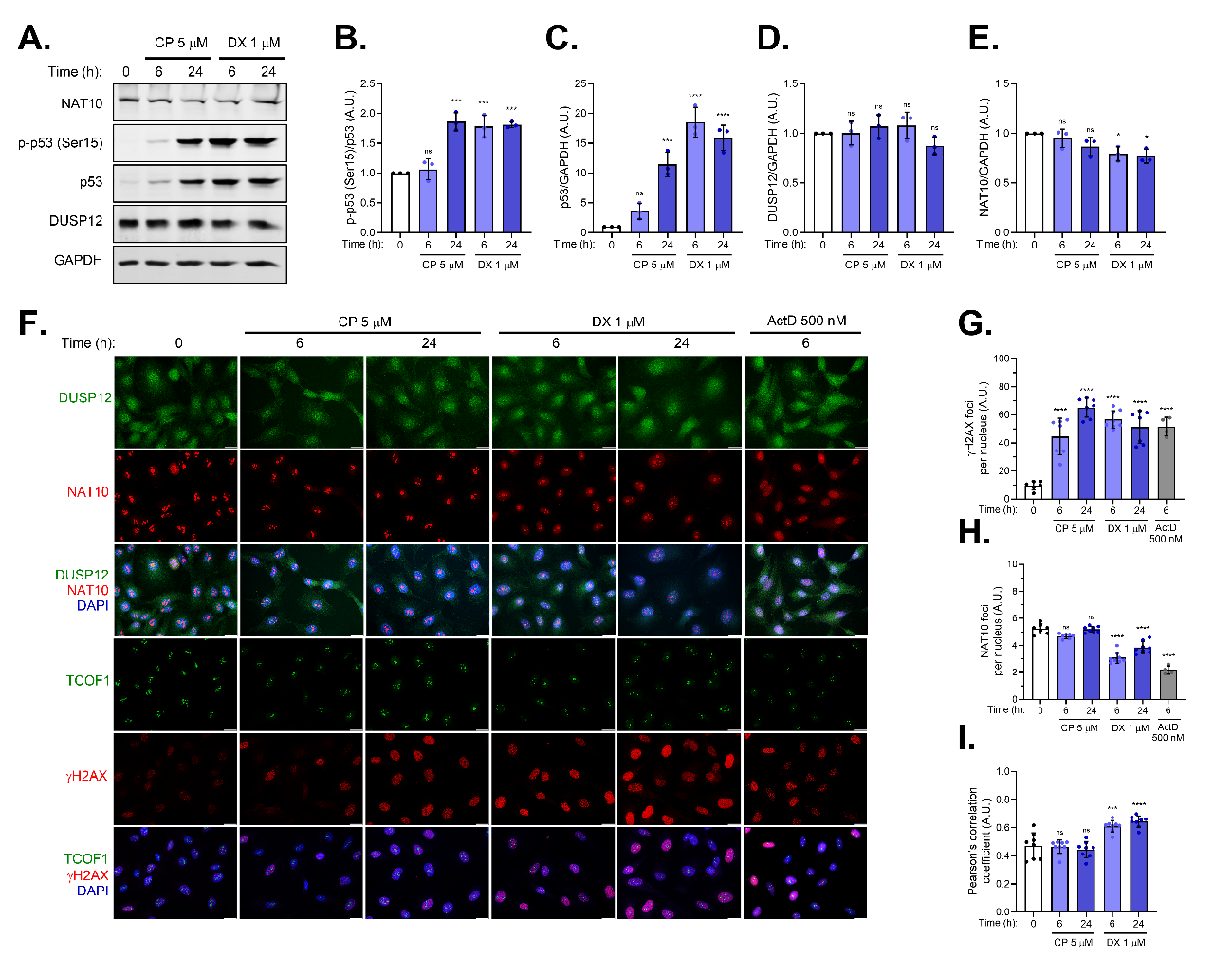

**Figure S3. DDR activation and NAT10 redistribution in HepaRG cells after genotoxic treatments.** (A) Western blot detection of NAT10, phospho-p53 (Ser15), total p53, DUSP12, and GAPDH in HepaRG cells treated with CP or DX. (B–E) Densitometric quantification of (B) phospho-p53 (Ser15), (C) total p53, (D) DUSP12, and (E) NAT10 relative to GAPDH expression. (F) Representative immunofluorescence images of DUSP12, NAT10, TCOF1, and γH2AX in HepaRG cells treated with CP, DX, or ActD. Scale bar = 20 μm. (G) Quantification of γH2AX foci per nucleus. (H) Quantification of NAT10 foci per nucleus. (I) Colocalization of DUSP12 and NAT10 by Pearson’s correlation coefficient. Data represent mean ± SD of at least three independent experiments. Statistical significance relative to control is indicated as *p < 0.05; **p < 0.01; ***p < 0.001; ****p < 0.0001, ns = not significant.

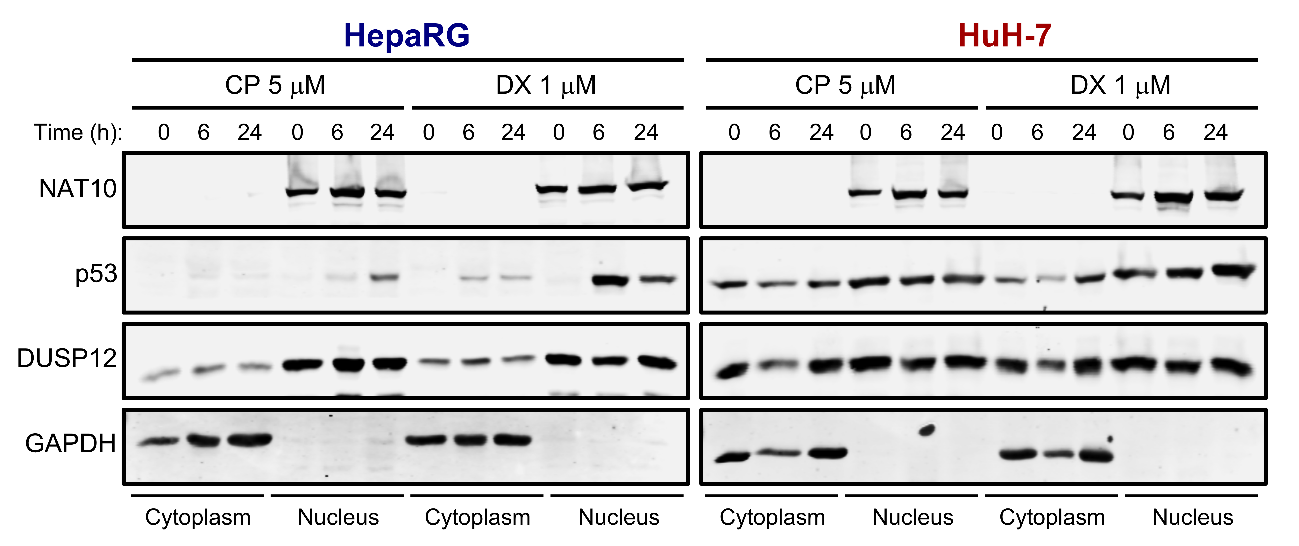

**Figure S4. Subcellular distribution of DUSP12 in HuH-7 and HepaRG cells after genotoxic treatments.** Western blot detection of cytoplasmic and nuclear fractions of HuH-7 and HepaRG cells treated with CP or DX. NAT10 and GAPDH served as nuclear and cytoplasmic fractionation controls, respectively; p53 was used as a marker of DDR activation.

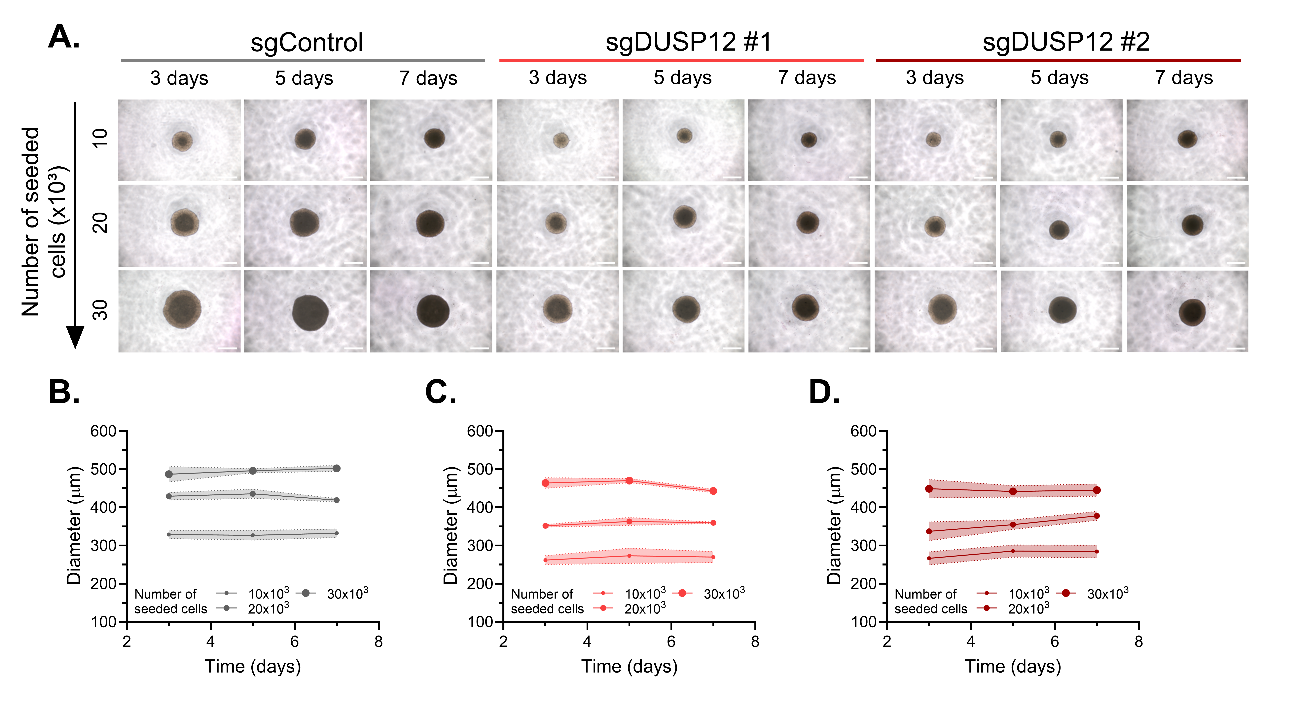

**Figure S5. Characterization of DUSP12 knockout clones growing in spheroids.** (A) Representative micrographs of DUSP12-KO HuH-7 spheroids generated by plating different initial cell numbers and maintained in culture for the indicated times. Scale bar = 200 µm. (B–D) Quantification of mean spheroid diameter (µm) of (B) sgControl, (C) sgDUSP12 #1, and (D) sgDUSP12 #2 clones under the same conditions. Graphs represent mean ± SD of at least three independent experiments.

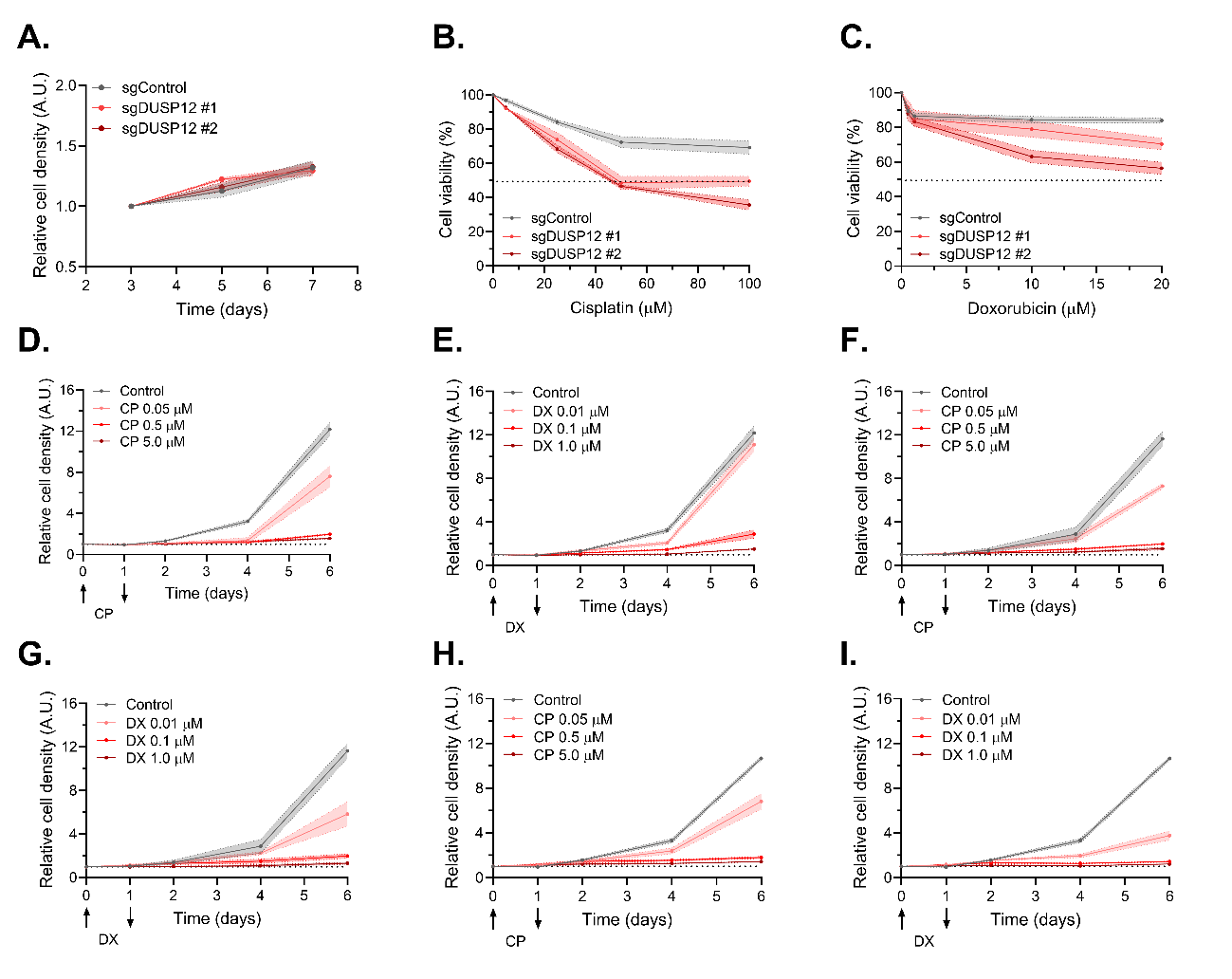

**Figure S6. Cytotoxic effects of cisplatin and doxorubicin in monolayer and spheroid cultures of DUSP12 knockout clones.** (A) Proliferation curves of DUSP12-KO HuH-7 spheroids. (B–C) Cell viability of DUSP12-KO HuH-7 clones in monolayer cultures treated with increasing concentrations of (B) CP or (C) DX for 72 h. (D–E) Proliferation curves of sgControl HuH-7 cells in monolayer cultures treated with increasing concentrations of (D) CP or (E) DX for 24 h. (F–G) Proliferation curves of sgDUSP12 #1 HuH-7 cells in monolayer cultures treated with increasing concentrations of (F) CP or (G) DX for 24 h. (H–I) Proliferation curves of sgDUSP12 #2 HuH-7 cells in monolayer cultures treated with increasing concentrations of (H) CP or (I) DX for 24 h. Graphs represent mean ± SD of at least three independent experiments.

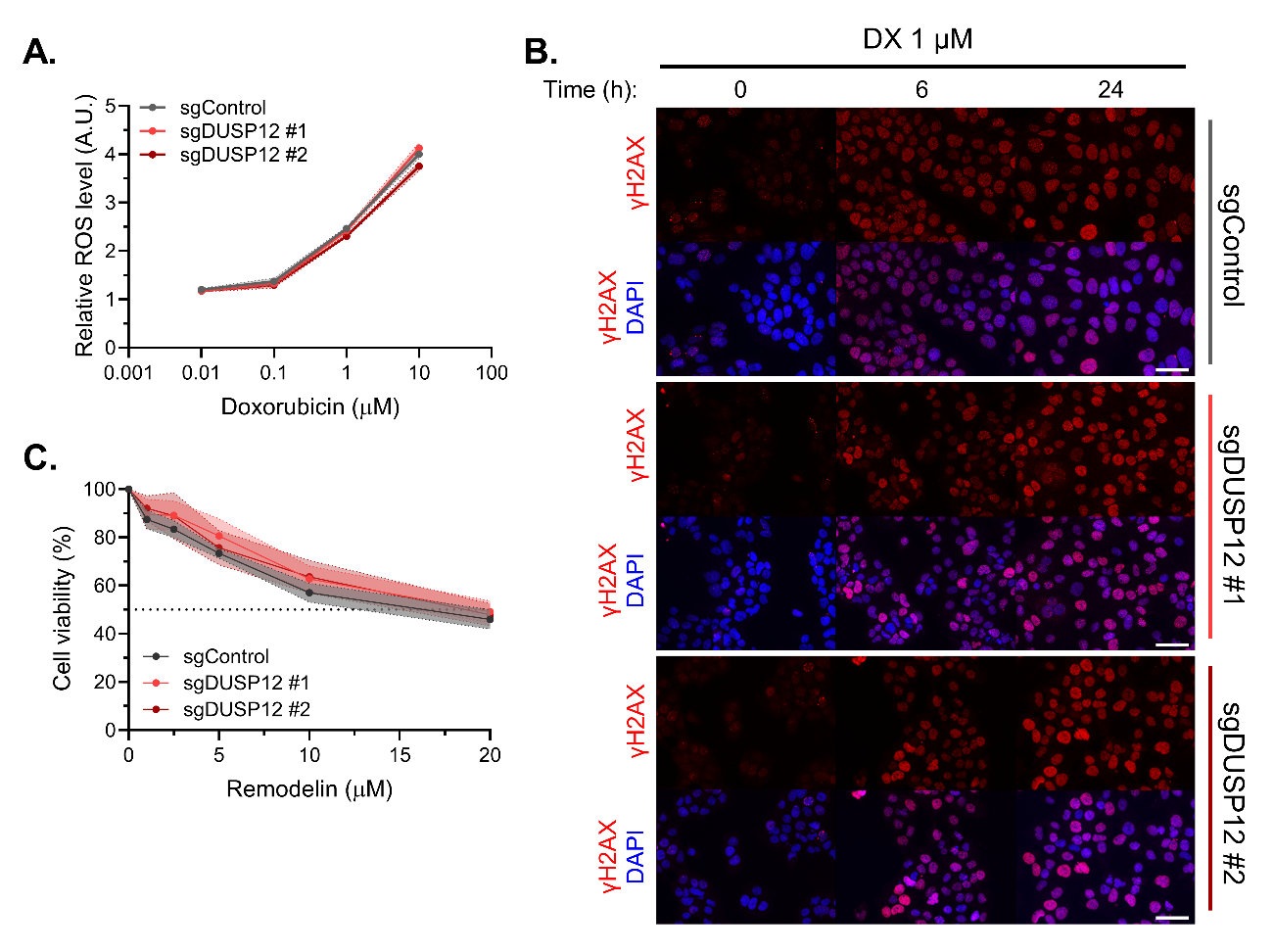

**Figure S7. Intracellular ROS levels and DDR activation in sgDUSP12 clones after doxorubicin treatments.** (A) Quantification of reactive oxygen species (ROS) levels in DUSP12-KO HuH-7 clones treated with increasing concentrations of DX for 6 h, relative to untreated controls. (B) Representative micrographs of γH2AX staining in DUSP12-KO HuH-7 clones treated with DX under the indicated conditions. Scale bar = 50 µm. (C) Cell viability of DUSP12-KO HuH-7 clones treated with increasing concentrations of RMD for 72 h. The graph represents the mean ± SD of at least three independent experiments.

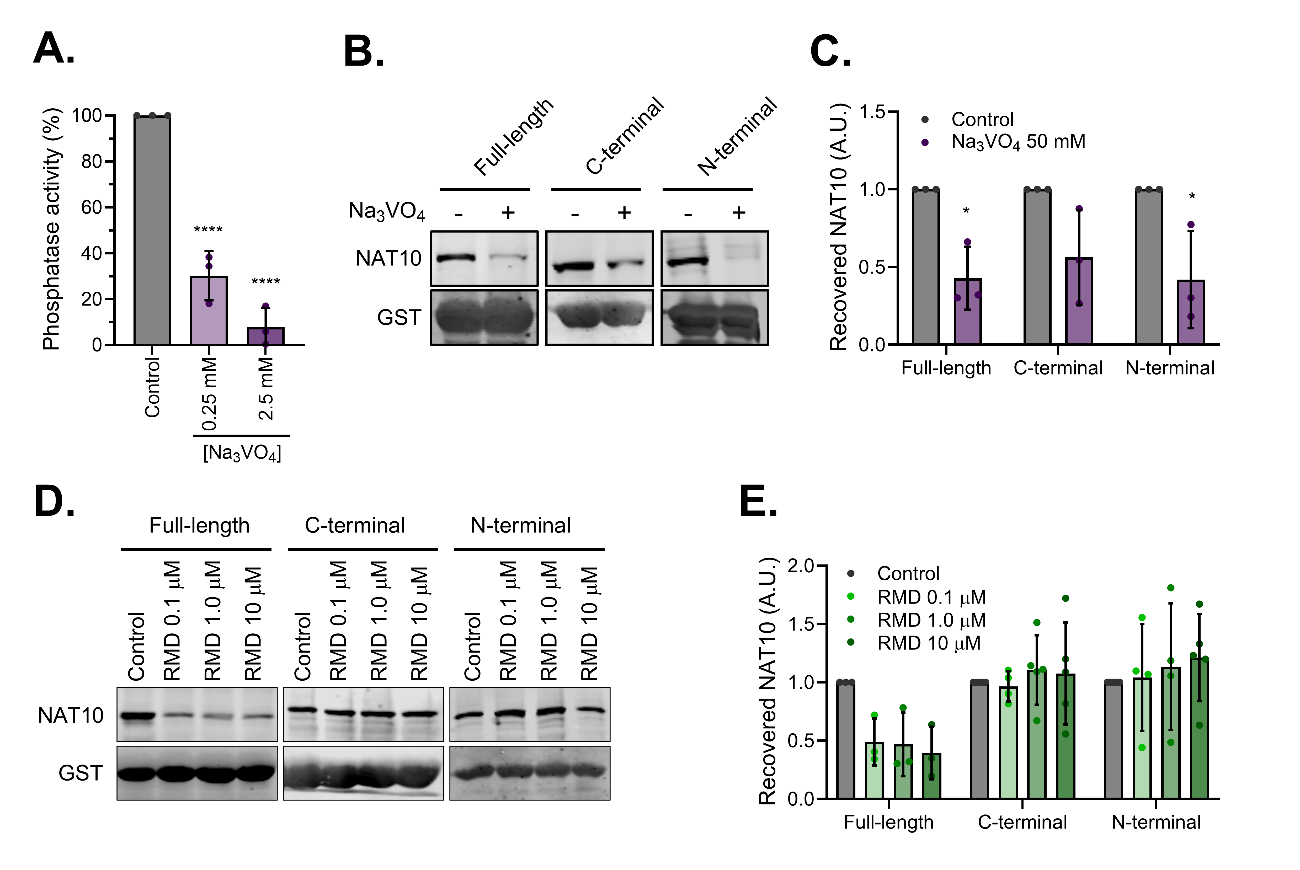

**Figure S8. DUSP12-NAT10 interaction is perturbed by catalytic activity inhibition of both proteins.** (A) Phosphatase activity of DUSP12 measured using 0.25 µg of purified protein in the presence of increasing concentrations of Na_3_VO_4_. Differences relative to control are indicated. Phosphatase activity was assessed using the artificial substrate DiFMUP (Invitrogen), with fluorescence monitored at 358/455 nm in the presence or absence of Na_3_VO_4_. (B) Western blot detection of NAT10 and GST after pull-down assays using 50 µg of purified DUSP12-GST constructs in the presence of 50 mM Na_3_VO_4_, adapted based on phosphatase inhibition observed at 250 µM Na_3_VO_4_. (C) Densitometric quantification of NAT10 levels after pull-down assays. Differences relative to control are indicated for each DUSP12 construct. (D) Western blot detection of NAT10 and GST after pull-down assays of NAT10 with different DUSP12-GST constructs in the presence of increasing concentrations of remodelin (RMD). (E) Densitometric quantification of NAT10 levels after pull-down assays. Statistical significance is indicated as *p < 0.05; **p < 0.01; ***p < 0.001; ****p < 0.0001, ns = not significant.

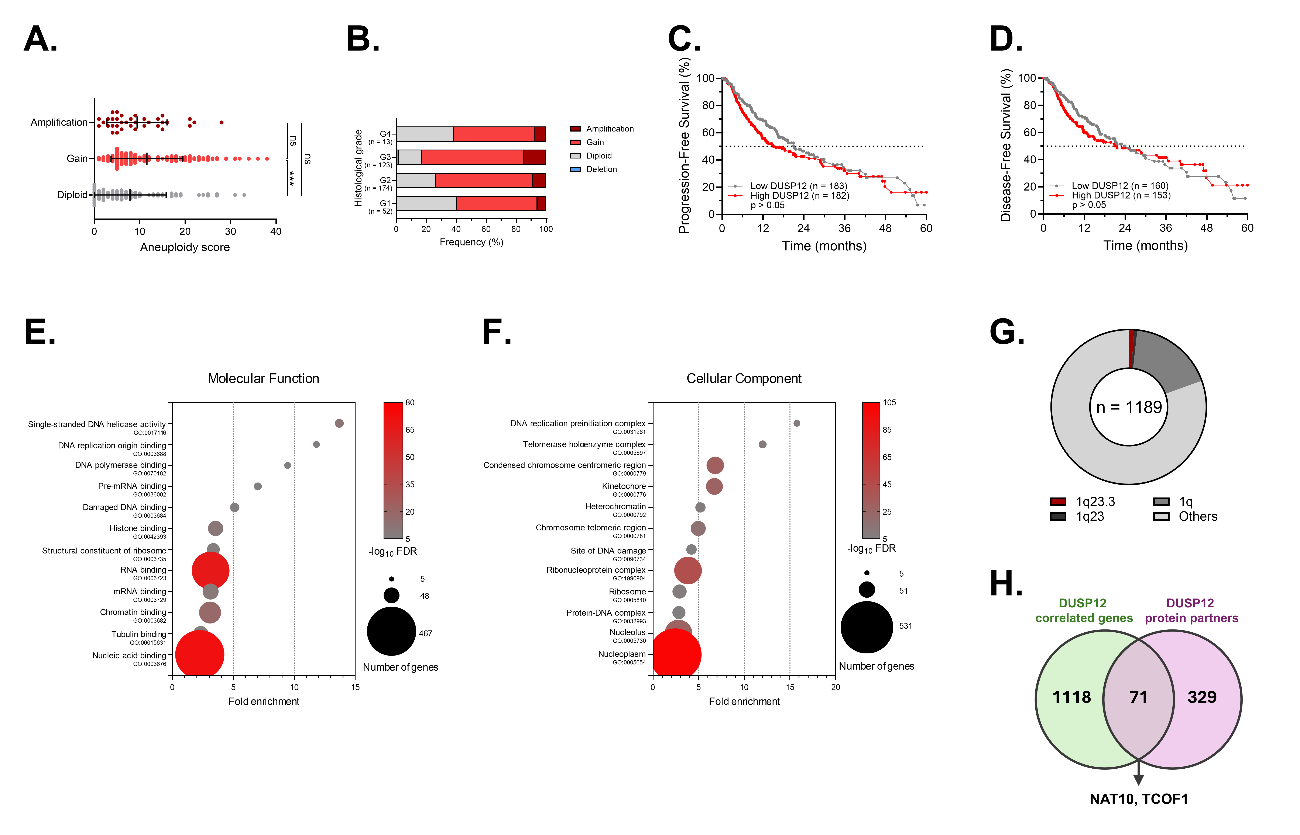

**Figure S9. Genomic instability and gene ontology enrichment associated with DUSP12 in HCC.** (A) Aneuploidy score in HCC patients with or without *dusp12* copy number alterations. (B) Distribution of DUSP12 copy number alterations in HCC patients according to histological grade. (C–D) Kaplan–Meier analyses of (C) progression-free survival and (D) disease-free survival at 5 years in HCC patients stratified by median DUSP12 expression. (E–F) Gene ontology analysis of (E) molecular function and (F) cellular component associated with genes correlated with DUSP12 expression in HCC patients. (G) Cytogenetic band distribution of genes correlated with DUSP12 expression in HCC patients used for gene ontology analyses. (H) Venn diagram showing the overlap between genes correlated with DUSP12 expression in HCC patients used for gene ontology analyses and reported protein partners of DUSP12.

**Supplementary Tables**

**Table S1.** Sequences of sgRNAs #1 and #2 used for the CRISPR knockout of DUSP12.

| **sgRNA** | **Guide sequence** |
| --- | --- |
| sgDUSP12 #1 | 5’–TGCAGGAGTCAGTCGAAGTG–3’ |
| sgDUSP12 #2 | 5’–GGATGAATGAGGGGTTTGAG–3’ |

**Table S2.** List of antibodies used for immunofluorescence assays.

| **Antibody** | **Manufacturer** | **Catalog number** | **Dilution** |
| --- | --- | --- | --- |
| DUSP12 | Santa Cruz Biotechnology | sc-390760 | 1:300 |
| NAT10 | Proteintech | 13365-1-AP | 1:500 |
| TCOF1 | Santa Cruz Biotechnology | sc-374536 | 1:300 |
| γH2AX | Cell Signaling Technology | #2577 | 1:500 |
| BrdU | Santa Cruz Biotechnology | sc-32323 | 1:300 |
| Geminin | Cell Signaling Technology | #52508 | 1:300 |
| Alexa Fluor-488 | Invitrogen | A11008, AA11001 | 1:500 |
| Alexa Fluor-568 | Invitrogen | A10042, A10037 | 1:500 |
| Alexa Fluor-647 | Invitrogen | A31573, A31571 | 1:500 |

**Table S3.** List of antibodies used for immunoblotting assays.

| **Antibody** | **Manufacturer** | **Catalog number** | **Dilution** |
| --- | --- | --- | --- |
| DUSP12 | Santa Cruz Biotechnology | sc-390760 | 1:2000 |
| NAT10 | Proteintech | 13365-1-AP | 1:5000 |
| p-p53 (Ser15) | Cell Signaling Technology | #9284 | 1:1000 |
| p53 | Santa Cruz Biotechnology | sc-56180 | 1:1000 |
| GAPDH | Cell Signaling Technology | #5174 | 1:2000 |
| β-actin | Invitrogen | PA1-183 | 1:4000 |
| ac4C | Proteintech | 68498-1-Ig | 1:1000 |
| GST | Invitrogen | XG345725 | 1:1000 |
| p-tyrosine | Sigma | P3300 | 1:2000 |
| Alexa Fluor-680 | Invitrogen | A21109, A21057 | 1:15000 |
| Alexa Fluor-800 | Invitrogen | A32735, A32730 | 1:15000 |

**Table S4.** IC_50_ values, coefficient of determination (R²), and 95% confidence interval (95% CI) from nonlinear regression of dose-response curves in HepaRG and HuH-7 in monolayer culture treated with cisplatin (CP) or doxorubicin (DX) for 48 or 72 hours.

| **HepaRG** | | | |
| --- | --- | --- | --- |
|  | | **48 h** | **72 h** |
| **CP** | IC_50_ (µM) | 16.4 | 12.4 |
|  | R² | 0.982 | 0.991 |
|  | 95% CI | 14.9 – 18.1 | 11.6 – 13.4 |
| **DX** | IC_50_ (µM) | 1.11 | 0.245 |
|  | R² | 0.822 | 0.922 |
|  | 95% CI | 0.635 – 1.88 | 0.169 – 0.332 |
| **HuH-7** | | | |
|  | | **48 h** | **72 h** |
| **CP** | IC_50_ (µM) | 14.9 | 8.10 |
|  | R² | 0.966 | 0.977 |
|  | 95% CI | 12.7 – 17.8 | 7.04 – 9.30 |
| **DX** | IC_50_ (µM) | 1.36 | 0.351 |
|  | R² | 0.968 | 0.986 |
|  | 95% CI | 1.11 – 1.66 | 0.306 – 0.399 |

**Table S5.** Doubling time values and 95% confidence interval (95% CI) from nonlinear regression of proliferation curves in HepaRG and HuH-7 in monolayer cultures treated with different concentrations of cisplatin (CP) or doxorubicin (DX) for 24 hours.

| **HepaRG** | | | | | |
| --- | --- | --- | --- | --- | --- |
| **CP** | **Concentration (µM)** | **0** | **1** | **5** | **25** |
|  | Doubling time (days) | 1.57 | 2.00 | 3.74 | 18.7 |
|  | 95% CI | 1.51 – 1.64 | 1.94 – 2.08 | 3.24 – 4.56 | 8.96 – ND |
| **DX** | **Concentration (µM)** | **0** | **0.5** | **1** | **5** |
|  | Doubling time (days) | 1.57 | 8.85 | 14.2 | 18.1 |
|  | 95% CI | 1.51 – 1.64 | 6.00 – 21.7 | 8.09 – 19.7 | 7.52 – 47.3 |
| **HuH-7** | | | | | |
| **CP** | **Concentration (µM)** | **0** | **1** | **5** | **25** |
|  | Doubling time (days) | 1.78 | 2.97 | 3.44 | 11.1 |
|  | 95% CI | 1.72 – 1.86 | 2.59 – 3.65 | 2.93 – 4.94 | 7.30 – 30.1 |
| **DX** | **Concentration (µM)** | **0** | **0.5** | **1** | **5** |
|  | Doubling time (days) | 1.78 | 5.41 | 6.11 | 7.37 |
|  | 95% CI | 1.72 – 1.86 | 4.12 – 8.98 | 4.60 – 10.3 | 5.54 – 12.2 |

**Table S6.** IC_50_ values, coefficient of determination (R²), and 95% confidence interval (95% CI) from nonlinear regression of dose-response curves in HepaRG and HuH-7 in spheroids treated with cisplatin (CP) or doxorubicin (DX) for 48 or 72 hours.

| **HepaRG** | | | |
| --- | --- | --- | --- |
|  | | **48 h** | **72 h** |
| **CP** | IC_50_ (µM) | 93.5 | 63.9 |
|  | R² | 0.969 | 0.958 |
|  | 95% CI | 83.1 – 108.7 | 56.6 – 73.1 |
| **DX** | IC_50_ (µM) | 26.9 | 9.63 |
|  | R² | 0.976 | 0.977 |
|  | 95% CI | 21.6 – 35.1 | 8.08 – 11.6 |
| **HuH-7** | | | |
|  | | **48 h** | **72 h** |
| **CP** | IC_50_ (µM) | 75.3 | 40.1 |
|  | R² | 0.936 | 0.905 |
|  | 95% CI | 61.3 – 98.0 | 30.8 – 52.8 |
| **DX** | IC_50_ (µM) | 71.9 | 52.9 |
|  | R² | 0.928 | 0.915 |
|  | 95% CI | 44.1 – 149 | 32.8 – 110 |

**Table S7.** IC_50_ values, coefficient of determination (R²), and 95% confidence interval (95% CI) from nonlinear regression of dose-response curves in DUSP12-KO clones in monolayer cultures treated with cisplatin (CP) or doxorubicin (DX) for 48 or 72 hours.

|  | | **sgControl** | **sgDUSP12 #1** | **sgDUSP12 #2** |
| --- | --- | --- | --- | --- |
| **CP** | IC_50_ (µM) | 28.9 | 37.4 | 24.9 |
|  | R² | 0.948 | 0.934 | 0.932 |
|  | 95% CI | 24.2 – 35.5 | 30.6 – 48.2 | 20.4 – 31.3 |
| **DX** | IC_50_ (µM) | 4.61 | 2.34 | 0.642 |
|  | R² | 0.980 | 0.940 | 0.974 |
|  | 95% CI | 3.99 – 5.34 | 1.82 – 3.04 | 0.534 – 0.763 |

**Table S8.** IC_50_ values, coefficient of determination (R²), and 95% confidence interval (95% CI) from nonlinear regression of dose-response curves in DUSP12-KO clones in spheroids treated with cisplatin (CP) or doxorubicin (DX) for 48 or 72 hours.

|  | | **sgControl** | **sgDUSP12 #1** | **sgDUSP12 #2** |
| --- | --- | --- | --- | --- |
| **CP** | IC_50_ (µM) | 276.2 | 74.4 | 50.5 |
|  | R² | 0.900 | 0.880 | 0.982 |
|  | 95% CI | 180.9 – 559.7 | 56.3 – 1105 | 45.9 – 55.9 |
| **DX** | IC_50_ (µM) | >1000 | 326.6 | 33.9 |
|  | R² | 0.555 | 0.770 | 0.956 |
|  | 95% CI | ND | 93.2 – ND | 24.8 – 51.3 |

**Table S9.** Doubling time values and 95% confidence interval (95% CI) from nonlinear regression of proliferation curves in DUSP12-KO clones in monolayer cultures treated with different concentrations of cisplatin (CP) or doxorubicin (DX) for 24 hours.

| **sgControl** | | | | | |
| --- | --- | --- | --- | --- | --- |
| **CP** | **Concentration (µM)** | **0** | **0.05** | **0.5** | **5** |
|  | Doubling time (days) | 1.70 | 2.18 | 7.09 | 10.2 |
|  | 95% CI | 1.63 – 1.78 | 1.98 – 2.50 | 6.02 – 8.82 | 8.75 – 12.4 |
| **DX** | **Concentration (µM)** | **0** | **0.01** | **0.1** | **1** |
|  | Doubling time (days) | 1.70 | 1.78 | 4.28 | 12.9 |
|  | 95% CI | 1.63 – 1.78 | 1.69 – 1.93 | 3.79 – 5.00 | 9.90 – 19.3 |
| **sgDUSP12 #1** | | | | | |
| **CP** | **Concentration (µM)** | **0** | **0.05** | **0.5** | **5** |
|  | Doubling time (days) | 1.74 | 2.16 | 6.20 | 9.95 |
|  | 95% CI | 1.65 – 1.84 | 2.05 – 2.30 | 5.92 – 6.52 | 8.96 – 11.3 |
| **DX** | **Concentration (µM)** | **0** | **0.01** | **0.1** | **1** |
|  | Doubling time (days) | 1.74 | 2.44 | 6.14 | 17.9 |
|  | 95% CI | 1.65 – 1.84 | 2.27 – 2.67 | 5.59 – 6.84 | 14.2 – 24.6 |
| **sgDUSP12 #2** | | | | | |
| **CP** | **Concentration (µM)** | **0** | **0.05** | **0.5** | **5** |
|  | Doubling time (days) | 1.79 | 2.24 | 6.59 | 10.9 |
|  | 95% CI | 1.73 – 1.87 | 2.11 – 2.40 | 5.93 – 7.48 | 9.69 – 12.7 |
| **DX** | **Concentration (µM)** | **0** | **0.01** | **0.1** | **1** |
|  | Doubling time (days) | 1.79 | 3.25 | 10.6 | 22.9 |
|  | 95% CI | 1.73 – 1.87 | 3.06 – 3.49 | 8.93 – 13.2 | 18.6 – 30.4 |

**Table S10.** IC_50_ values, coefficient of determination (R²), and 95% confidence interval (95% CI) from nonlinear regression of dose-response curves in DUSP12-KO clones pre-treated with 10 µM RMD for 24 h prior to exposure to cisplatin (CP) or doxorubicin (DX) for 48 h (in the presence of RMD).

|  | | **sgControl** | **sgDUSP12 #1** | **sgDUSP12 #2** |
| --- | --- | --- | --- | --- |
| **CP** | IC_50_ (µM) | 32.5 | 39.6 | 34.3 |
|  | R² | 0.954 | 0.946 | 0.932 |
|  | 95% CI | 28.7 – 37.3 | 34.6 – 46.5 | 29.4 – 41.1 |
| **CP + RMD** | IC_50_ (µM) | 31.3 | 40.3 | 35.6 |
|  | R² | 0.933 | 0.939 | 0.923 |
|  | 95% CI | 26.4 – 38.3 | 34.6 – 48.4 | 29.4 – 45.1 |
| **DX** | IC_50_ (µM) | 4.19 | 2.75 | 0.668 |
|  | R² | 0.966 | 0.977 | 0.972 |
|  | 95% CI | 3.60 – 4.93 | 2.43 – 3.13 | 0.573 – 0.775 |
| **DX + RMD** | IC_50_ (µM) | 2.79 | 0.928 | 0.305 |
|  | R² | 0.973 | 0.994 | 0.966 |
|  | 95% CI | 2.43 – 3.22 | 0.867 – 0.994 | 0.254 – 0.362 |

**Table S11.** IC_50_ values, coefficient of determination (R²), and 95% confidence interval (95% CI) from nonlinear regression of dose-response curves in DUSP12-KO clones treated with remodelin (RMD) for 72 h.

|  | | **sgControl** | **sgDUSP12 #1** | **sgDUSP12 #2** |
| --- | --- | --- | --- | --- |
| **RMD** | IC_50_ (µM) | 15.9 | 19.1 | 18.2 |
|  | R² | 0.948 | 0.880 | 0.876 |
|  | 95% CI | 13.9 – 18.8 | 15.6 – 25.4 | 14.7 – 24.1 |
